# Neo-sex Chromosomes Anchor a Sex-Limited Polymorphism Under Gene Flow

**DOI:** 10.64898/2026.06.28.735067

**Authors:** P. L. H. de Mello, J. K. Kelly, R. E. Glor

## Abstract

How polymorphic traits are maintained despite the homogenizing forces of gene flow and recombination is a central question in evolutionary biology. While the accumulation of locally adaptive alleles within chromosomal inversions is well established, the role of sex chromosomes in local adaptation has received comparatively less empirical support. Here, we show that a male-limited color polymorphism in an *Anolis distichus* contact zone is anchored by a neo-sex chromosome. Across a narrow environmental gradient, an abrupt transition between yellow and orange dewlaps—extensible throat fans used for signaling—is driven by distinct pigmentation profiles we characterize via histological and chromatographic analyses. Integrating genomics and association mapping, we demonstrate this divergence relies on an additive, oligogenic architecture. Alternative neo-Y haplotypes track the phenotypic cline and combine additively with autosomal variants near a putative ketolase and a lipid regulator to determine color, jointly explaining a significant portion of the phenotypic variance. Furthermore, we identify a copy number variant near an X-linked visual processing gene, indicating simultaneous sensory divergence. This concerted evolution suggests the local light environment drives adaptation of the communication system. Ultimately, this study demonstrates that degrading neo-sex chromosomes act as non-recombining hubs for locally adaptive alleles, preserving phenotypic diversity under gene flow.

## Main

Understanding how phenotypic diversity arises and persists is a central goal of evolutionary biology. Sexually dimorphic color patterns represent one of the most spectacular manifestations of this diversity. Because sexually dimorphic traits impact fitness through natural and sexual selection, they serve as powerful models for studying the interplay of ecology, behavior, and genomic architecture^1,2^. Among sexually dimorphic traits, sex-limited polymorphisms—exclusive to the heterogametic sex—must decouple from the shared genome to resolve inter-sexual conflict while maintaining intra-sexual variation against the homogenizing forces of gene flow and recombination^3^. To protect co-adapted alleles against the homogenizing forces of gene flow, loci under divergent selection are predicted to accumulate in regions of reduced recombination, like inversions^4^. Heteromorphic sex chromosomes not only exhibit reduced recombination relative to autosomes, but also house a disproportionate number of genes underlying sexual conflict and are influenced by demographic factors like sex-biased dispersal. These dynamics amplify allele frequency divergence between the X and Y, making sex chromosomes a compelling hub for loci under sexual and natural selection^5–7^. Despite a strong theoretical basis, empirical support remains scarce for the role of sex chromosomes in establishing sexual dimorphism and local adaptation^8–10^, in part due to the inherent challenges of performing phenotype-to-genotype associations for sex-linked traits.

While unchallenged gene flow erodes genetic divergence and hampers local adaptation, stable contact zones across environmental clines capture the balance of selection, dispersal, and local adaptation, making them natural laboratories for genetic inference^11–13^. These processes are expected to produce a largely homogeneous genome where regions associated with the trait under selection show the highest differentiation^14^. Furthermore, within the core of the contact zone multiple generations of admixture allow recombination to erode genome-wide linkage disequilibrium (LD)^15^. This decay of background genetic structure isolates the targets of selection, facilitating high-resolution genotype-phenotype mapping. Therefore, a system where polymorphism in a sexually dimorphic trait tracks an environmental cline offers an unique opportunity to test the hypothesis that sex chromosomes play a disproportional role in local adaptation. Such a system exists in the remarkably diverse neotropical anole lizards.

Anole lizards are a well-established system for studying ecology, evolution and behavior, propelled into the ‘omics era by a wealth of recent genomic resources^16–18^ (Fig. 1). One of the most striking features of anoles is the extensible, often brightly colored throat fan, or dewlap, that males display during highly stereotyped signaling behaviors^19^. The dewlap is thought to play a central role in species recognition and sexual selection, as its coloration and patterning are generally conserved within species. One notable exception is a widespread species, the Hispaniolan Bark anole–*Anolis disitchus*,– occurring across Hispaniola, its satellite islands, and the Bahamas that exhibits striking geographic variation in dewlap coloration and pattern, ranging from pale yellow to deep wine red^20^. Because many of these forms exchange genes where they come into contact, this variation is generally interpreted as subspecific differentiation rather than strict reproductive isolation, and may instead reflect local adaptation to signaling conditions, with yellow dewlaps favored in drier environments and orange or red dewlaps favored in wetter environments.

**Figure 1:**
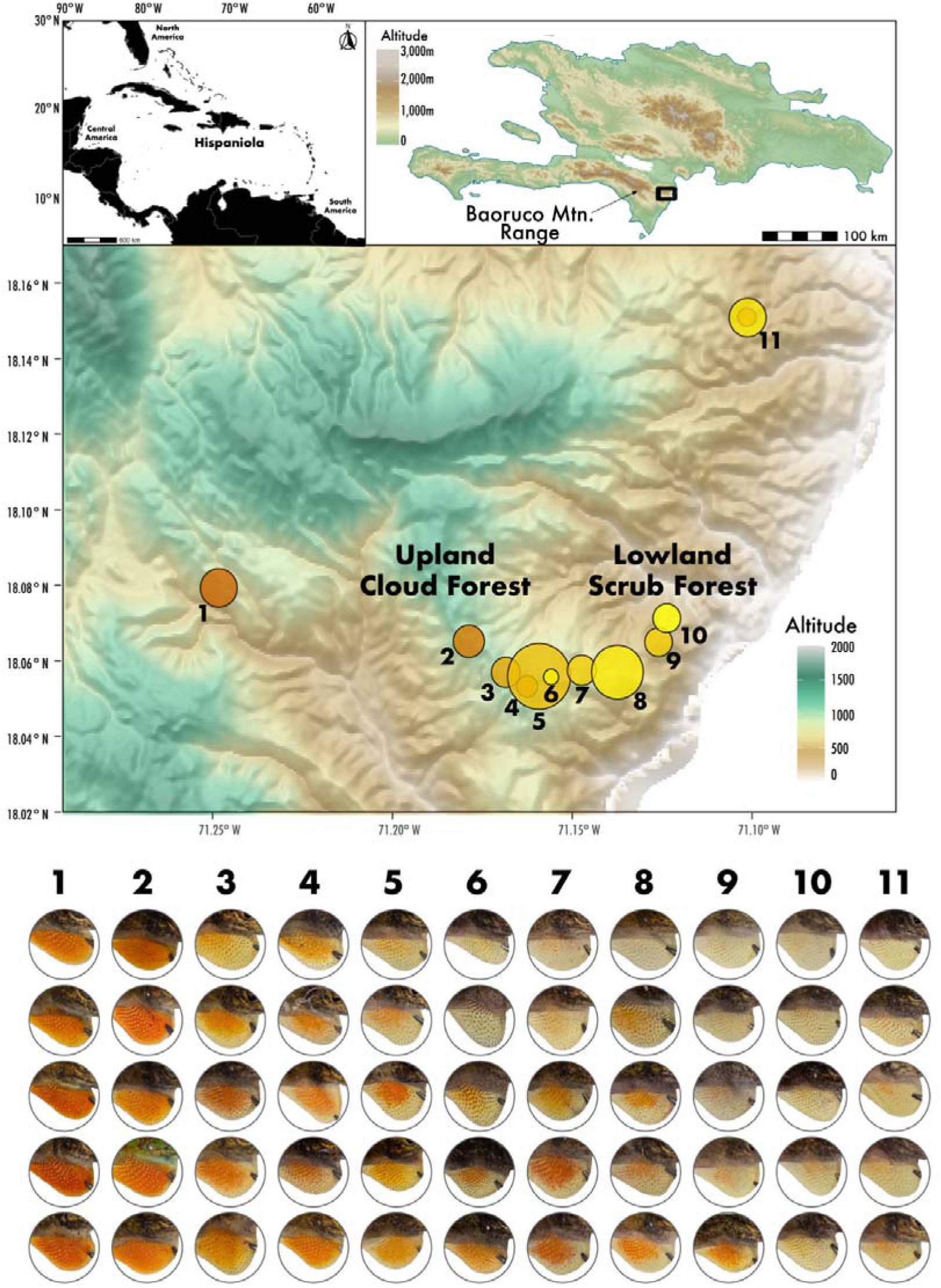
Sampling and phenotypic variation of *Anolis distichus favillarum* from the southern Dominican Republic. (a) The Island of Hispaniola. (b) Sampling area in the Baoruco Mountains. (c) Geographic transect, with sampling effort represented by circle radius and mean hue by color. (d) Representative dewlap phenotypes from each locality.

Our prior work has shown that dewlap color in the Hispaniolan bark anole dewlap is highly heritable and oligogenic^21,22^. Some phenotypically distinct populations show genetic differentiation and reduced gene flow where they come into contact, suggesting that they may represent incipient species. However, other populations exhibit abrupt phenotypic transitions without detectable genetic differentiation. A prominent example occurs in the Bahoruco Mountains of the southern Dominican Republic, where decades of work using classic genetic markers such as allozymes and mtDNA have found no evidence of genetic differentiation despite a striking shift in dewlap coloration from yellow in dry coastal forest to orange in mesic montane forest within a single recognized subspecies–*Anolis distichus favillarum*^23^.

Furthermore, we recently assembled the *A. distichus* genome, as well as its XXY neo-sex chromosome system, confirming this system was formed through multiple autosomal fusions to the ancestral X and Y chromosomes shared across all Iguanians^18^. Importantly, if a formerly autosomal neo-Y chromosome starts to degrade post-fusion, then we expect these fusions to capture standing genetic variation previously segregating within autosomes in environments that cease to recombine with their neo-X homolog. Consequently, neo-sex chromosomes are uniquely positioned to, like inversions, act as “supergenes” during divergence with gene flow^24^. By uniting a sharp phenotypic cline and unconstrained gene flow in a diverse group with neo-sex chromosomes, the *A. d. favillarum* contact zone offers an unparalleled system to investigate the genomic architecture of local adaptation.

We integrate population genomics, pigment biochemistry, and skin histology, finding compelling support for a disproportionate role of neo-sex chromosomes in maintaining dewlap color polymorphism in *A. distichus* across a phenotypic cline. First, we demonstrate that a neo-Y-linked haplotype strictly tracks the phenotypic cline, resisting introgression from migrating males. Second, we show that this neo-Y-haplotype anchors an oligogenic, additive network, interacting with autosomal *TMEM68* (Transmembrane Protein 68) and *CYP2J2-like* loci to additively explain 13.3% of the phenotypic variance —providing further evidence that *CYP2J2-like* (Cytochrome P450 Family 2 Subfamily J Member 2-like) acts as the primary carotenoid ketolase in squamates^25,26^. In addition, we identify an outstanding difference in allele frequency between orange– and yellow-dewlapped specimens for an X-linked cone arrestin (*ARR3*, Arrestin 3), a gene that mediates visual adaptation to the local photic environment^27^. This results provides evidence that both signal (color) and receptor (vision) are co-evolving across the environmental transect. Ultimately, our results illustrate that an oligogenic system anchored on a heteromorphic neo-Y chromosome combines additively with autosomes to maintain a sexually dimorphic polymorphism in the face of gene flow.

### The Hemizygous *A. distichus* XXY System

To test whether sex chromosomes contribute to dewlap color variation, we characterized the degradation of neo-Y-linked regions in the XXY system of *A. distichus*^6^. First, we used whole-genome resequencing (WGS) of males and females to generate high-resolution maps of female-to-male read depth ratios, sex-specific heterozygosity, and SNP density^28^. During this process, we identified and corrected a chimeric assembly artifact within scaffold 12, the Neo-X/Y pair in *A. distichus* (see Supplementary Methods; Supplementary Fig. 1). Our genomic characterization of the *A. distichus* XXY system with WGS data revealed extensive Y degradation and X hemizygosity in males (Fig. 2). As expected for hemizygous regions, X-linked scaffolds averaged a log2(F:M) depth ratio of ∼1 alongside elevated female heterozygosity^28^, while neo-Y-linked scaffolds showed deeply male-biased coverage (log2(F:M) < –1; Fig. 2a).

**Figure 2:**
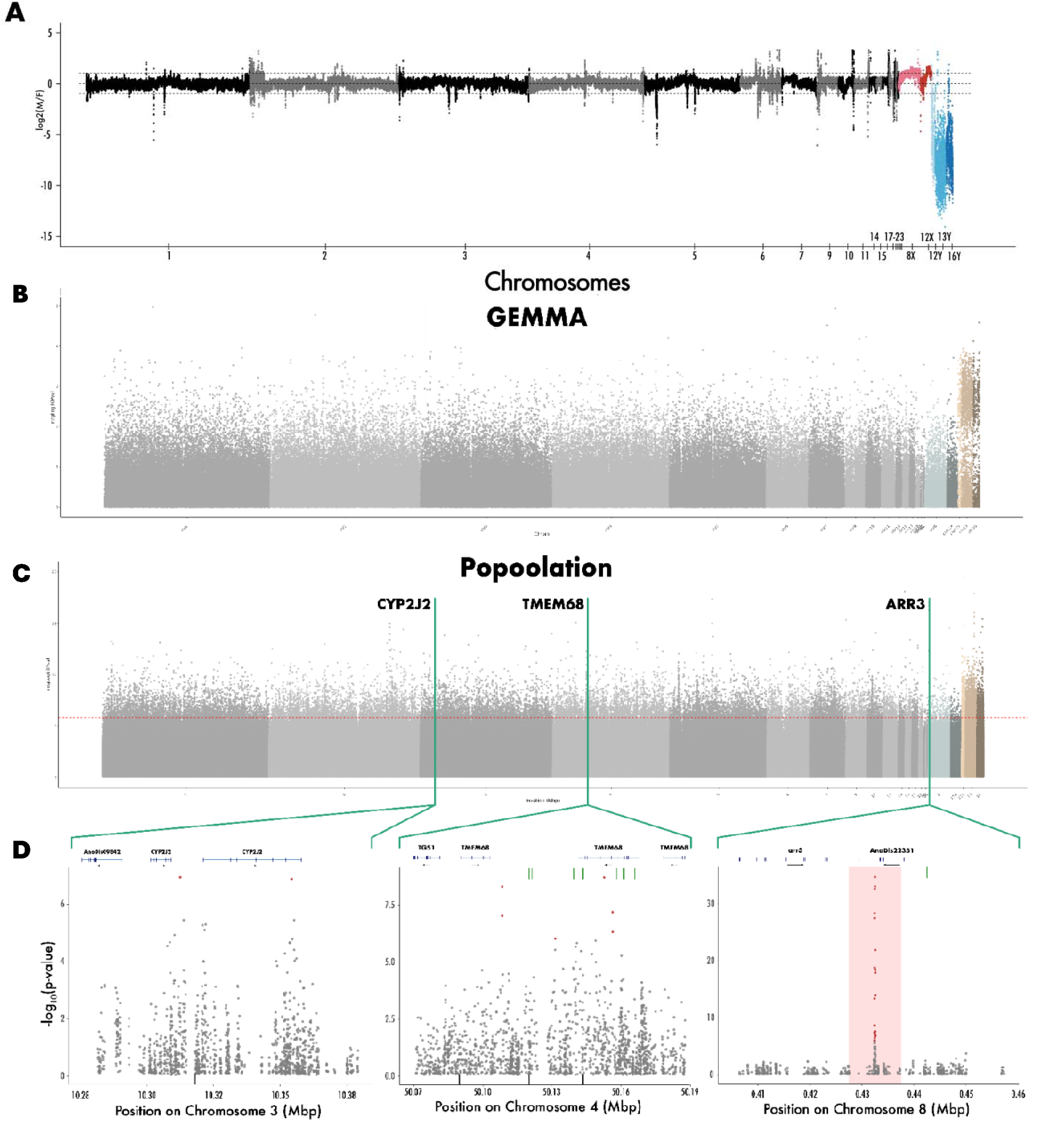
Sex chromosome hemizygosity in the *A. distichus* XXY system and genotype-phenotype association Manhattan plots showing oligogenic Neo-Y-linked and autosomal basis of color pattern. (a) Sliding-window analysis (10 kbp) of weighted female-to-male log2 read depth. Dashed lines represent autosomal (0), X-linked (∼1), and degraded Y-linked (≤ –1) expectations. Autosomes (gray), X-linked (red), and Y-linked scaffolds (blue) are ordered numerically. (b) Manhattan plot of GWA statistics for dewlap hue; red line denotes the permutation-based significance threshold. (c) –log10(p-value) from QTLSeqR estimated through *G’* statistics between orange and yellow pools (dashed line = 99th percentile). (d) Magnified regions of interest containing *CYP2J2-like* (left) and *TMEM68*(center) and *ARR3* (right) showing Fisher’s Exact Test values (black dots) and MSG markers (red triangles). Regions flagged as containing CNVs are highlighted by a red-shaded rectangle.

Sliding window analyses confirmed thorough degradation across most of the neo-Y, despite the relative recency of the autosome-to-sex-chromosome fusions that produced it^29^. Two regions, however, escaped this degradation. First, a terminal region of scaffold 8—the ‘ancient X,’ —which retains autosomal depth patterns (Fig. 2c) with few sex specific variants and slightly reduced coverage (Supplementary Fig. 2). Given the acrocentric morphology of most *Anolis* microchromosomes^29^, this likely corresponds to a shared centromeric region between the ancient X and Y. Second, a region of scaffold 12X which displays autosomal depth patterns and equal male-female heterozygosity, characterizing it as a non-degraded pseudoautosomal region (Supplementary Fig. 1; Fig. 2b). We corroborated these assignments using population-level linkage disequilibrium (LD) from our reduced representation ‘GWA dataset’ (see below). As expected, LD decayed fastest on autosomes and slowest on non-recombining neo-Y-linked scaffolds (Supplementary Fig. 3). Together these results confirm a predominantly hemizygous system, enabling a direct test of how neo-sex chromosome dynamics contribute to the maintenance of a sex-limited polymorphism.

## Cellular and Pigmentary Basis of Color

A detailed characterization of the phenotype is essential for evolutionary interpretation of genetic mapping results. To this end, we investigated the biochemical and cellular basis of the color phenotype. Based on prior research in anoles^30^, we hypothesized that the dewlap color polymorphism in *Anolis distichus favillarum* reflects quantitative and qualitative differences in pigment composition, specifically dietary carotenoids and pteridines. To test this, we sampled 11 male *A. d. favillarum* across a micro-transect (< 2 km) that captured the full range of color phenotypes without showing any evidence for genome-wide population structure (Fig. 1a).

Spectrophotometric analysis of sequentially extracted carotenoids and pteridines confirmed that dewlap color variation corresponds to both quantitative differences in carotenoid concentration and qualitative differences in pteridine composition (Fig. 3a). Spectrophotometric analysis revealed a dual pigmented system. Green backs and orange dewlaps show a three-peaked absorbance spectrum characteristic of carotenoids that were absent in yellow dewlaps (*p* < 0.001; Tukey HSD). Orange dewlaps also showed higher maximum absorbance than yellow dewlaps (*p* < 0.001; Welch two sample T-test), indicating a higher concentration of molecules within that absorbance spectrum in orange than yellow dewlaps (Supplementary Fig. 4). Our prior transcriptomic work, for example, found that *SCARB1* (Scavenger receptor class B type 1) —a transmembrane carotenoid transporter that is X-linked in anole lizards—is upregulated in orange relative to yellow skin in a second subspecies pair of Orange / Yellow dewlap populations within *A. distichus*^26^. Additionally, orange dewlaps were the only tissues exhibiting an absorbance spectrum characteristic of red pteridines (presumably drosopterins), indicating the existence of a dual-pigment system in *A. distichus* analogous to the vibrant reds seen in the closely related *Anolis sagrei*^30^.

**Figure 3:**
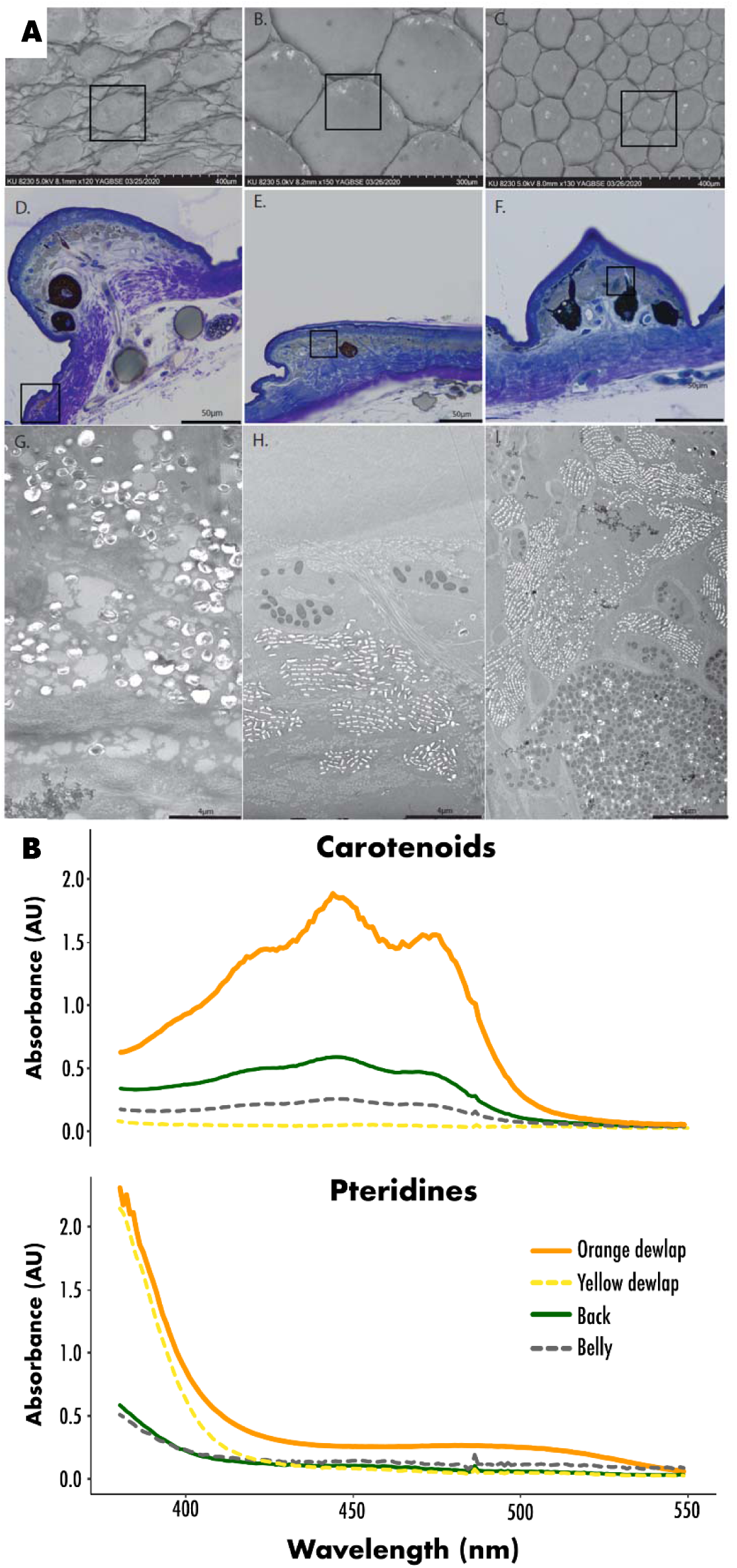
Cellular and biochemical analyses reveal the structural and pigmentary basis of dewlap hue. (a) Scanning (top), light (middle), and transmission electron microscopy (TEM; bottom) show dewlap hinges contain a single layer of xanthophores packed with carotenoid and pteridine vesicles. Dewlap hinge skin notably lacks the iridophores and melanophores present in belly and back skin. (b) Light absorbance spectra for orange dewlap, yellow dewlap, back, and belly tissues. Carotenoids form the three-peaked distribution (peak ∼446 nm), while the lower absorbance peak (450–500 nm) unique to the orange dewlap indicates drosopterin-like pteridines.

Because pigment distribution is regulated by cellular architecture, we next used light microscopy, scanning electron microscopy (SEM), and transmission electron microscopy (TEM) to examine how these biochemical differences map onto skin histology. Ultrastructural analysis revealed that unlike the multi-layered ‘chromatophore unit’ present in scales–with ‘color cells’ organized in layers– the extensible dewlap folds possess a simplified structure (Fig. 3b): a single dense layer of yellow/orange xanthophores, with black melanophores and shiny iridophores effectively absent. Within this xanthophore layer, cells are packed with two distinct organelle types—large lipid-like vesicles, which house apolar carotenoids, and smaller structures containing concentric rings, which house polar pteridines. The absence of underlying iridophores and melanophores in the dewlap folds suggests that dewlap color is a direct readout of xanthophore pigment content, unmodulated by structural color or melanin-based interference. This simplified, direct relationship between both pigment biochemistry and visual phenotype allows us to link genetic variation to color divergence across the contact zone.

## Neo-Y Resists Genome-Wide Introgression

To test the prediction that neo-sex chromosomes anchor the divergence of this sexually dimorphic signal, we examined the spatial distribution of allele frequencies across the contact zone. We sampled the genome of 287 specimens distributed across 11 localities (Fig. 1a) with genome-wide restriction-site associated markers (Multiplexed Shotgun Genotyping loci, MSG loci) and fit geographic cline models to the phenotypic data (dewlap hue measured from RAW photographs) and each MSG locus. We calculated allele frequency changes for 794,802 loci, and found only 13,484 (1.69%) to have an allele frequency change of at least 30% across the cline, confirming an extensively shared genomic background between orange and yellow dewlapped *A. d. favillarum*^23^. Consistent with our prediction, clinal analysis identified a disproportionate number of candidate loci on sex-limited scaffolds relative to its length, specifically on the non-recombining neo-Y-linked scaffolds (# neo-y-linked = 1,017; # non-y-linked = 12,467, p-value Binomial Test < 2.2e-16; Fig. 4).

**Figure 4:**
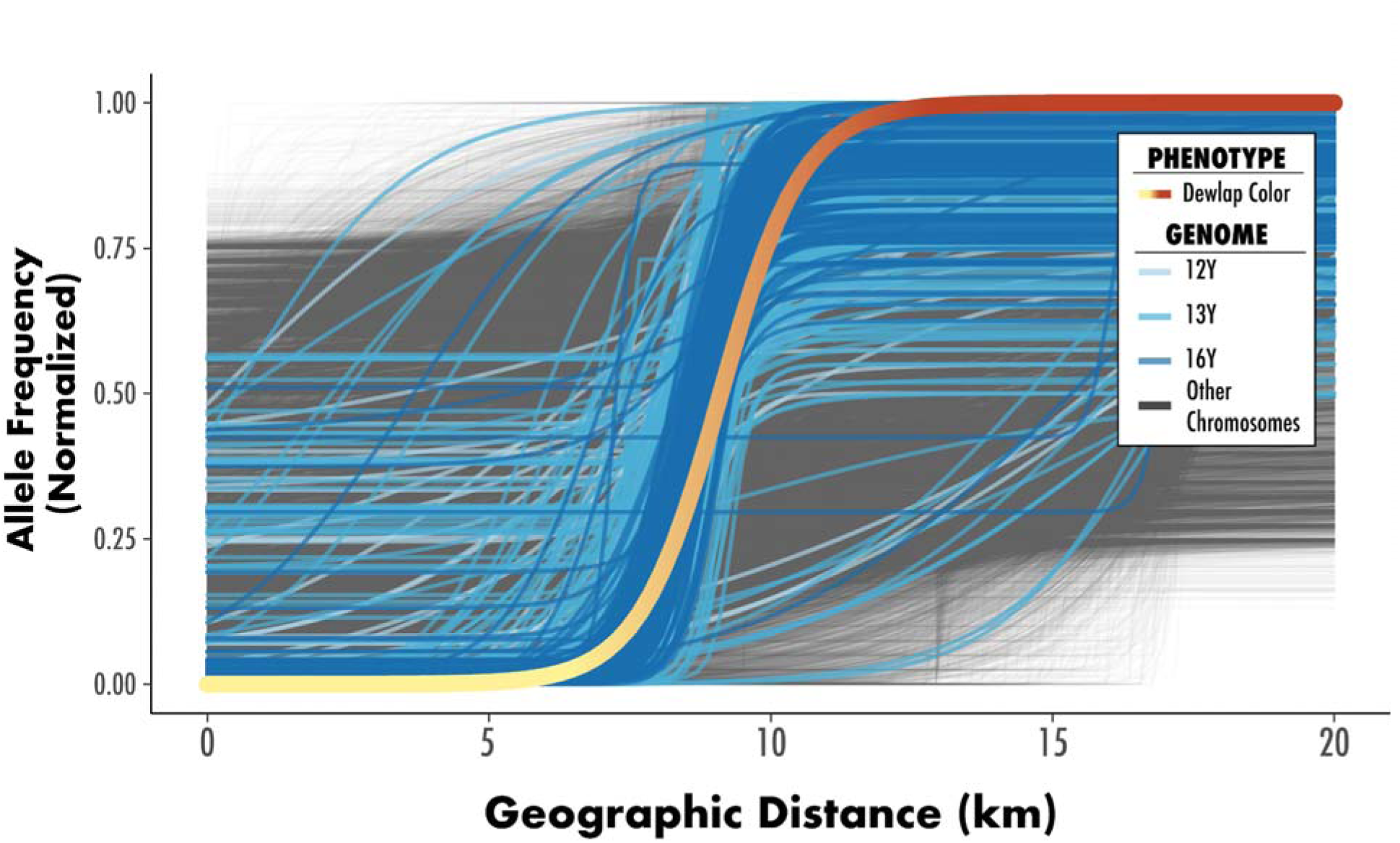
Neo-Y-linked allele frequencies strongly track clinal variation in dewlap color. Normalized, ascending clines for loci with >30% allele frequency divergence across the geographic transect. Scaled phenotypic values for dewlap hue (yellow to orange) are overlaid to highlight the strict spatial concordance between the phenotypic transition and neo-Y-linked loci clines (blue shades).

Because the neo-Y-linked scaffolds of *A. distichus* are extensively degraded (Fig. 2a), we expected recombination rates would be lower between neo-Y-linked loci than between X-linked or autosomal loci. Accordingly, we observed long distance linkage disequilibrium (LD) across all three neo-Y-linked scaffolds (Supplementary Fig. 3). The reduced recombination of the Y allowed us to phase neo-Y-linked variants into two discrete haplotypes segregating across the contact zone. A Principal Components Analysis (PCA) including all neo-Y-linked loci split the specimens into two non-overlapping haplotypic groups (Hap0 and Hap1), with PC1 explaining over 50% of the variance. Fitting a cline model according to neo-Y-haplotype assignment recovered a steep transition, with each haplotype fixed at either extreme of the transect. Furthermore, the neo-Y-haplotype cline closely tracked both the slope and center of the dewlap color cline (Supplementary Fig. 5).

Theory predicts that loci under divergent selection with gene flow will accumulate in regions of reduced recombination to prevent the breakup of co-adapted alleles; the *A. d. favillarum* degrading neo-Y chromosome fulfills this requirement by sheltering these loci in a large, non-recombining haplotypic block^32^. Our results are consistent with the prediction that neo-sex chromosomes are uniquely positioned to harbor such blocks—even if transiently^24^—and that neo-sex-linked sequences may contribute disproportionately to adaptive divergence under gene flow^5^.

## Neo-Y Linkage to Color Variation

Having established that the neo-Y chromosome is under strong divergent selection across the contact zone, we next sought to explicitly map the genetic basis of dewlap color. To confirm the mechanistic link between neo-Y-linked sequences and the male-limited phenotype, we conducted a Genome-Wide Association (GWA) analysis in populations at the center of our cline, testing for associations between MSG loci and dewlap color. We restricted this analysis to 195 specimens collected from the center of the contact zone (localities 5-8 in Fig. 1) to minimize the impact of background population structure (Supplementary Fig. 6), the main source of false positives in GWA analyses.

We imputed missing genotypes, and tested for genotype-to-phenotype associations using linear mixed models (LMM) while accounting for population stratification. The LMM model confirmed the clinal findings: neo-Y-linked loci are strongly associated with the yellow-to-orange dewlap color polymorphism (Fig. 5b). We further evaluated the genetic architecture of dewlap color using a Bayesian Sparse Linear Mixed Model (BSLMM). While extensive linkage disequilibrium within the neo-Y-haplotype prevented the identification of a single causal locus via BSLMM with high posterior inclusion probability (PIP, the Bayesian estimate of a variant’s independent causal effect)^33^ variants on neo-Y-linked scaffolds exhibited almost 10 fold larger sparse effect sizes (β, the magnitude and direction of the effect) than autosomal or X-linked variants, highlighting the disproportionate contribution of neo-Y-linked variants (non-Y mean β = 1.243; non-Y mean standard error = 0.008; y mean β = 11.72; y mean standard error = 0.2779712; Wilcoxon rank sum test, *p* < 2.2e-16; Supplementary Fig. 7). This concordance between spatial introgression and localized genotype-phenotype association provides compelling evidence linking the non-recombining neo-Y chromosome and the implementation of this sex-limited polymorphism.

**Figure 5:**
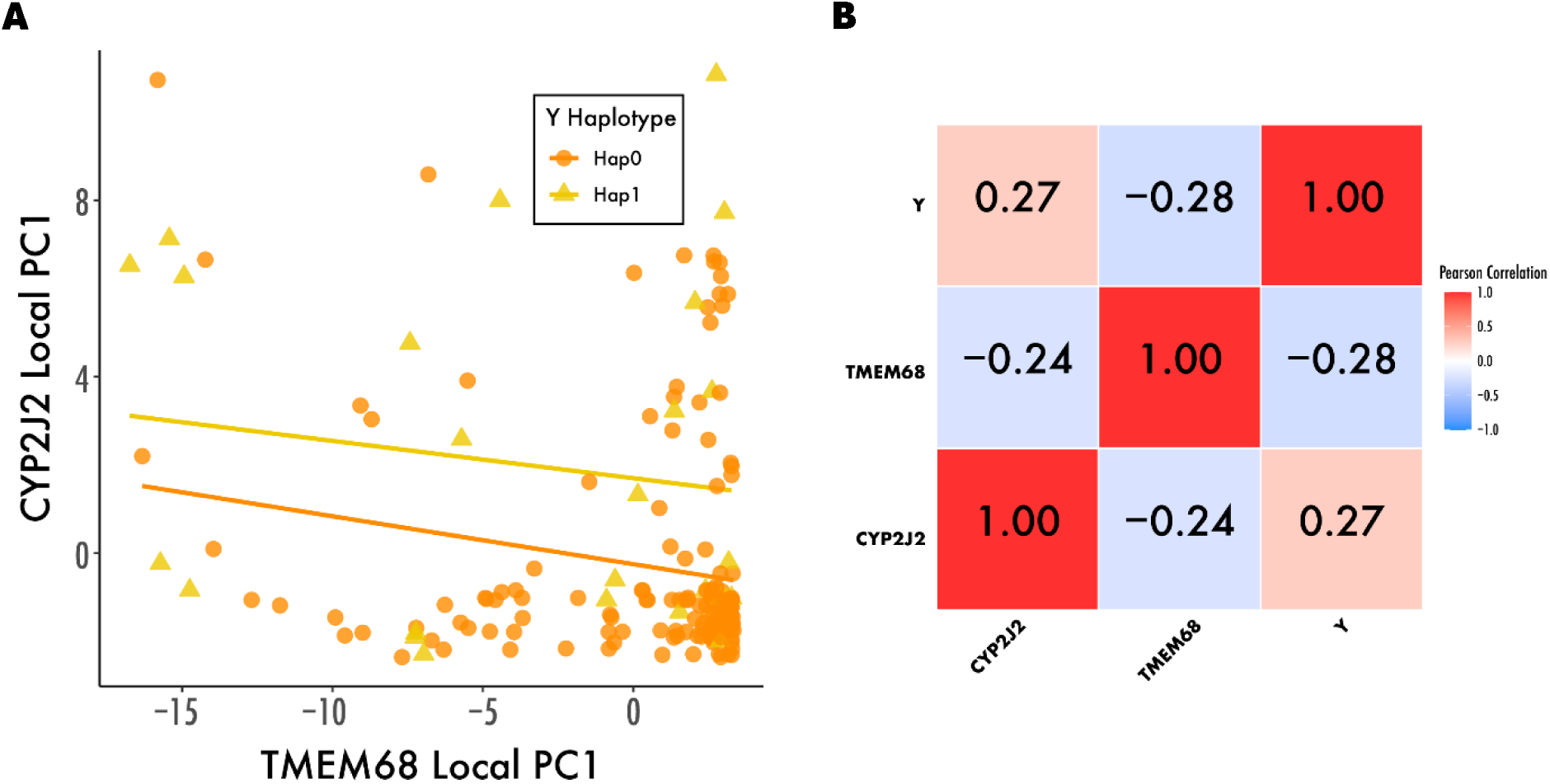
Inter-chromosomal linkage disequilibrium maintains co-adapted allele networks for dewlap hue. (a) Genotypic distribution based on the first principal component (PC1) of variants spanning *CYP2J2-like* and *TMEM68*, colored by neo-Y-haplotype. (b) Correlation matrix testing gametic phase disequilibrium among the three predictive components of the joint additive model. Significant inter-chromosomal correlations indicate non-random assortment among these loci.

## BSA Confirms Autosomal Candidate Genes

Because rapid LD decay limits the resolution of MSG data (Supplementary Fig. 3), we fine-mapped candidate loci using bulk segregant analysis (BSA) through whole-genome pooled sequencing on 50 of the most orange and 50 of the most yellow specimens from the contact zone (Supplementary Fig. 6). After testing for genotype-phenotype associations, we cross-referenced BSA variants with GWA and clinal loci. We considered variants as candidates if they met one of two criteria: (i) a significant BSA association (*q* ≤ 0.05) within 25 kbp of a GWA (*p* ≤ 0.05) or clinal locus, or (ii) a top 0.01% BSA association for variants located beyond the 25-kbp window. To contextualize these candidates, we cross-referenced them against known vertebrate color-patterning genes and transcripts differentially expressed between orange and yellow dewlaps in *A. distichus*^26^.

Criterium (i) identified candidate variants not only in neo-Y-linked regions, but also at autosomal regions in the vicinity of *CYP2J2-like* and *TMEM68* gene clusters (Supplementary Data 1; Fig. 5; see below). The *CYP2J2-like* gene cluster is a known mediator of carotenoid ketolation^34^ we previously identified as differentially expressed between orange and yellow dewlaps in *A. distichus*^26^. *TMEM68*, on the other hand, has been linked to polymorphisms in coat color in cattle^35^, that plays an important role in cellular lipid storage and membrane lipid homeostasis^36^ and has not yet been associated with orange/yellow polymorphisms. Both regions encompass localized clusters of paralogous genes, a genomic architecture well-documented as an engine of evolutionary novelty that allows for tissue-specific specialization^37^. This tandem duplication is highly reminiscent of the *CYP2J2-like* clusters in zebra finches, where variation in copy number predicts red versus yellow beak morphs^34^. Reinforcing their putative causal role, multiple variants within these two autosomal clusters ranked in the top 1% of the most strongly associated loci to phenotype genome-wide.

Criterium (ii) revealed a highly significant region on the ‘ancient X’ chromosome (Fig. 5) that is located immediately downstream of *ARR3*. *ARR3* is crucial for quenching the phototransduction cascade in cone cells and optimizing visual processing^38^. Interestingly, *ARR3* overlaps with a region of the ancient X exhibiting pseudo-autosomal patterns of coverage (Supplementary Fig. 2), thus implying that this vision-related gene is present in two copies across both sexes. Because non-coding structural variants frequently drive rapid adaptation by altering *cis*-regulatory enhancer-promoter interactions^39^, we investigated the structural landscape of this downstream *ARR3* peak. We confirmed that this outlier window overlaps with a Copy Number Variant (CNV) segregating between the orange and yellow pools (Fig. 5). The presence of a divergent structural variant capable of altering the *cis*-regulation of a critical vision gene strongly suggests that local selection is optimizing not just the visual signal, but the sensory reception system itself across the environmental gradient^40^. Together, the neo-Y-linked haplotype, the autosomal pigment regulatory network (*CYP2J2-like* and *TMEM68*), and the X-linked visual modulator (*ARR3*) reveal a comprehensive genomic architecture of signal-receiver coevolution. The full list of candidate loci can be found at Supplementary Data 1.

## Neo-Y Anchors an Additive Color Network

Although our clinal and association data identify the neo-Y chromosome as the largest driver of phenotypic divergence, the exact molecular mechanism remains elusive. To identify putative candidates for the color polymorphism we then further annotated neo-Y-linked scaffolds. As expected, the annotation of the non-recombining neo-Y-linked scaffolds resulted in the identification of genes associated with sex determination in diverse vertebrate systems (*MAP3K4* [Mitogen-Activated Protein Kinase Kinase Kinase 4] and *DNMT3B* [DNA Methyltransferase 3 Beta] on scaffold 13; *SOX3* [SRY-Box Transcription Factor 3] on scaffold 16) alongside one gene that may play a role in the synthesis of guanine platelets in iridophores (*PRPSAP2* [Phosphoribosyl Pyrophosphate Synthetase Associated Protein 2] on scaffold 13^41^) and one gene that may act in the fate determination of undifferentiated chromatophore precursors by interacting with *SOX10* [SRY-Box Transcription Factor 10] (*SOX3* on scaffold 16^42^; Supplementary Data 2-4).

The lack of a candidate with direct effect on carotenoid deposition/modification or chromatophore fate in the neo-Y has led us to hypothesize that the neo-Y haplotype influences color polymorphism through an uncharacterized upstream mechanism. Accordingly, models of sex-limited polymorphisms have assumed the necessity of epistatic “master switches” to prevent the expression of unfit intermediate phenotypes^43^. To test this, we evaluated epistatic interactions between neo-Y haplotypes and our autosomal candidate loci (*CYP2J2*-like and *TMEM68*). Fitting linear models to the primary axes of genetic variation (Local PC1) across these candidate regions refuted the epistatic model (Fig. 6). Instead, an additive model between the neo-Y haplotype and autosomes best fit the data, with the major-effect neo-Y-linked haplotype explaining 5.29% (*p* < 0.001) of the variance in dewlap hue, while the autosomal loci *CYP2J2-like* and *TMEM68* explaining an additional 5.06% (*p* < 0.001) and 2.95% (*p* < 0.011) of the variance, respectively. This model is also consistent with the phenotypic data in the field (Fig. 1), where the neo-Y haplotype is sufficient but not necessary to explain part of the variance in dewlap color. Rather, it operates additively with autosomal loci to determine the final color pattern. Verifying this trans-regulatory network requires targeted *in vivo* assays, therefore continuing to develop such functional genomic tools in squamates represents the critical next frontier for understanding complex trait architecture.

## Consequences of Neo-Sex Linkage

The spectacular phenotypic diversity in systems like anole lizards is often attributed to ecological opportunity^44^, yet the genetic architectures protecting this diversity from the homogenizing forces of gene flow and recombination remain obscure. We hypothesized that the *A. distichus* dewlap polymorphism is maintained by strong directional selection anchored by its unique neo-sex chromosome configuration. Our results confirm this prediction. We observe steep, environment-tracking clines in both signal and receiver loci, alongside strong inter-chromosomal linkage disequilibrium (Pearson’s *r* ∼ 0.25) between unlinked autosomal and neo-Y-linked haplotypes at the center of the contact zone (Fig. 6). The lack of neo-Y-haplotype introgression and the simultaneous divergence of a vision-associated gene point to sensory drive, wherein strong selection acts against phenotypically mismatched males migrating across the environmental gradient.

The *Anolis distichus* dewlap color system provides a compelling contrast to classic models of local adaptation under gene flow. Evolutionary theory and empirical studies demonstrate that populations typically overcome the homogenizing effects of recombination either through the gradual expansion of linkage disequilibrium^14^ or, most commonly, through the “capture” of locally adaptive alleles within chromosomal inversions. However, there is growing theoretical and empirical evidence highlighting the disproportionate role of sex-linked loci in facilitating local adaptation^10^, and the sex-linkage of sexually dimorphic traits^45^. Accordingly, in the *A. d, favillarum* system divergence is anchored by a neo-Y-linked driver that acts in concert with at least two autosomal loci despite extensive gene flow. Our study is one of the few that empirically finds an association between sex chromosomes, let alone a neo-sex chromosome, and a locally adaptive polymorphism^46,47^.

We hypothesize that the autosome-to-sex-chromosome fusion in *A. distichus* acts in a similar fashion to inversions, effectively locking together locally adaptive combinations of loci^24^. By capturing tightly linked, locally adaptive loci and locking them within a region of suppressed recombination, this neo-sex linkage prevents the dissolution of multi-locus traits across the environmental gradient. With the advancement of second– and third-generation sequencing approaches, it has become clear that independent neo-sex fusions are frequent across *Anolis*^17,18^. Because each fusion incorporates different sets of autosomes, this structural turnover traps new combinations of loci within a non-recombining environment. Combined with subsequent neo-Y-chromosome degradation, this dynamic generates unique non-recombining hubs for locally adaptive alleles, equipping different clades with distinct genetic substrates for adaptation. As the discovery of neo-sex systems accelerates, it is increasingly apparent that chromosomal turnover is not merely a structural curiosity, but potentially a primary engine of phenotypic novelty and local adaptation.

## Materials and Methods

### Sampling & Phenotyping

We collected 287 adult male lizards from 11 localities across a 17.29 km transect located in the Baoruco Mountain range on the Barahona Peninsula in Southern Hispaniola (Fig. 1). We gathered phenotypic data by photographing each specimen using a Nikon D810 camera with a 105 mm macro lens (AF-S NIKKOR 105mm f/2.8 G Micro ED) and illumination by dual lens-mounted flashes (Nikon 4803 R1C1 Wireless Close-Up Speedlight Flash System Commander Kit). We photographed each specimen on white paper with a 1mm grid and included a Calibrite ColorChecker Passport in each photograph for standardization during post-processing. We developed custom *awk* and *R* code to extract RGB values from each RAW image, and followed Endler and Mielke^1^ to convert RGB values into brightness, chroma and hue (BCH) values. We quantified dewlap color as the average value for hue within a standardized 1×1 mm patch at the center of the dewlap, as this specific region best visually captured the phenotypic variation observed across the contact zone.

### Histology

We examined skin morphology and chromatophore ultrastructure of dewlap, belly, and back tissues using, Light Microscopy (LM), Scanning Electron Microscopy (SEM) and Transmission Electron Microscopy (TEM). For SEM, skin samples from one specimen were fixed, stained with osmium tetroxide, critical point dried, and imaged on a Hitachi SU8230. For TEM, samples (N=3 specimens) were fixed, post-fixed, stained with osmium and enhanced with p-phenylenediamine, dehydrated, embedded in Embed 812 resin, sectioned, stained with uranyl acetate and lead citrate, and imaged on a Hitachi H-8100 (Detailed methods in Supplementary Methods). We conducted all histological analyses at the University of Kansas’ Microscopy and Analytical Imaging (MAI) core.

### Pigment Biochemistry

We used absorbance spectroscopy to compare pigment content across dewlap, belly, and back skin. Pigments were sequentially extracted: first carotenoids using methanol:ethyl acetate (6:4, v/v) + BHT, then pteridines using 2% ammonium hydroxide following McLean et al.^2^. Absorbance spectra (UV-VIS-NIR) were measured for each extract relative to a blank. Samples exceeding 2 AU were serially diluted before measurement (Full details in Supplementary Methods).

### Library Preparation and Sequencing

We extracted high-molecular weight DNA from all samples using a custom bead-based protocol (modified from Rohland & Reich^3^). To characterize patterns of sex-chromosome degradation we performed whole-genome sequencing (WGS) on DNA extracted from two male and two female *A. distichus*. For the GWA and geographic cline analyses, we generated multiplexed shotgun genotyping (MSG) libraries following Adolfatto et al.^4^. For Bulk Segregant Analysis (BSA), we selected the 50 individuals with the most extreme orange and yellow dewlap hues from the GWA dataset, creating two pools, each with three independent technical replicates to minimize amplification bias^5^. WGS and BSA libraries were prepared using the Swift Ws Turbo DNA kit (Swift Biosciences) and sequenced by the Oklahoma Medical Research Facilities (OMRF) on an Illumina NovaSeq 4000 (150 bp paired-end). MSG libraries were sequenced in-house on an Illumina HiSeqX (150 bp paired-end). Target coverage was 10x for individual WGS samples, 10x for MSG samples, and 50x per pool (approx. 16.5x per replicate) for BSA.

### De-Multiplexing, QC, Alignment, Variant-Calling and Filtering

We processed raw MSG reads using *STACKS* ‘process_radtags’^6^ and WGS/BSA reads with *fastp*^7^. We aligned reads to the *A. distichus* genome using *BWA*^8^, followed by variant calling with *BCFtools* ‘mpileup’ and ‘call’^9^. The resulting variants were subsetted and filtered differently for each analysis (WGS, BSA, GWA, and Cline datasets) using custom scripts and *VCFtools*^10^. We applied custom thresholds for mapping quality, call quality, minor allele frequency, missing data, and Hardy-Weinberg equilibrium (details in Supplementary Methods). Missing genotypes in the MSG-derived GWA and Cline datasets were imputed using *BEAGLE*^11^.

### Characterizing Hemizygosity Across the Neo-Sex Chromosomes of *A. distichus*

To characterize the extent of hemizygosity and structural degradation across the *A. distichus* neo-sex chromosome complex, we analyzed whole-genome sequencing data from two males and two females. We calculated depth of coverage using *SAMtools*^12^ and extracted variant calls using *BCFtools*^10^. To standardize baseline coverage across specimens, we implemented custom *awk* scripts to compute the weighted mean depth per scaffold. We then calculated the log2-transformed ratio of female-to-male read depth [log2(F:M)] across 100 kbp sliding windows with 10 kbp overlapping steps.

Alongside read depth, we quantified the spatial distribution of genetic variation by calculating the proportion of segregating single nucleotide polymorphisms (SNPs) and sex-specific variants within the identical 100 kbp sliding windows, following^13^. These metrics evaluate the genomic signatures of recombination suppression^14^. Under this framework, diverging Y chromosomes accumulate male-specific SNPs due to strict sex linkage. As structural degradation progresses, the loss of neo-Y-linked sequence renders males hemizygous, resulting in a drop in male heterozygosity relative to females. These combined metrics partition the sex chromosomes into distinct evolutionary strata: hemizygous X-linked regions exhibit a 2:1 female-to-male depth ratio (log2[F:M] ≈ 1) coupled with female-biased heterozygosity; pseudo-autosomal regions (PARs) exhibit equal coverage (log2[F:M] ≈ 0) and shared variation; and non-recombining, neo-Y-linked regions display male-biased read depth (log2[F:M] < 0) accompanied by a high density of male-specific SNPs. We corroborated these findings by analyzing linkage disequilibrium (LD) patterns within the larger GWA dataset using *PLINK* v.1.9^15^ (--r2 –-ld-window-kb 1000000), expecting higher LD on non-recombining regions of Y scaffolds.

### Population Structure

We first verified the extent of genetic structure across the data by performing a Principal Component Analysis (PCA) with *PLINK* v.1.9. Prior to performing the PCA with default parameters, we used *PLINK* v.1.9 to filter unlinked variants in our Cline dataset with a maximum amount of missing data per locus of 20% and a maximum of 30% missing data per specimen. This dataset represented the best compromise between completeness of the dataset and number of loci sampled. Importantly, overall PC values did not vary qualitatively with the completeness of the matrix.

### Cline Analysis

We used R’s package *HZAR*^16^ to fit geographic clines across localities from the Barahona transect for both our phenotypic data and variants of our Cline dataset. Because the neo-Y chromosome extensively lacks recombination (see above), we hypothesized that it is inherited as a single unit, and that we should thus be able to phase neo-Y-linked reads into two or more haplotypes segregating across the transect. We then performed a Principal Component Analysis (PCA) on all neo-Y-linked variants to phase the non-recombining neo-Y-linked scaffolds (13, 16, and 12Y). As expected, the major axis of genetic variation (PC1) polarized the divergent neo-Y-haplotypes. Males were assigned to one of two discrete haplotype states based on the sign (positive or negative) of their individual PC1 score. These discrete assignments were then used to calculate locality-level haplotype frequencies, effectively formatting the entire neo-Y-chromosome as a single bi-allelic locus.

We modeled the spatial introgression of each autosomal, X-linked, and neo-Y-linked markers, as well as the composite neo-Y-haplotype. To thoroughly explore the cline’s shape, we fitted a comprehensive set of 15 models, representing all combinations of variance scaling (none, fixed, free) and exponential tails (none, left, right, both, mirror). Parameter estimation was performed using a two-step MCMC process. After a short initial run to find appropriate starting values, we initiated three independent, longer chains for each model, run for 1 x 10 generations with a 1 x 10 burn-in. To ensure robust exploration of the parameter space, starting values for key parameters were randomized for each replicate chain. The single best-fit model was selected based on the lowest Akaike Information Criterion corrected for small sample sizes (AICc), from which we extracted the maximum likelihood parameters for cline center and width. We also fitted a cline for dewlap hue, to assess the center and width of the phenotypic cline.

### Genome-Wide Association Analyses

We performed genotype-to-phenotype associations between imputed variants from the GWA dataset and dewlap hue using *GEMMA v0.98*^17^. We tested for locus-specific associations using a linear mixed model (-lmm4). The genetic relationship matrix (GRM) used by linear mixed models to account for genome-wide population structure can reduce statistical power if candidate loci are included in its calculation^18^. To prevent this proximal contamination—a phenomenon where the inclusion of causal loci in the kinship matrix causes the mixed model to inadvertently absorb the trait’s variance, attenuating true association signals^18^,— we calculated the GRM using autosomal and X-linked markers, excluding the neo-Y-linked scaffolds. To further model the hypothesized oligogenic architecture of the trait, we applied a Bayesian Sparse Linear Mixed Model (-bslmm 1) in *GEMMA* to extract variant-specific posterior inclusion probabilities (PIP) and sparse effect sizes (β).

### Bulk Segregant Analyses Fine Mapping

First, we calculated the Fixation Index (Fst) between orange and yellow pools in the BSA dataset across 20 kbp windows with 10 kbp steps using *grenedalf*^19^ with default parameters to identify regions of high divergence between pools. Then, to fine map the causal locus underlying the orange yellow polymorphism we utilized *QTLseqr*^20^ to identify variants from our BSA dataset that shows significant allele frequency differences between the orange and yellow pools. This method leverages local linkage disequilibrium to mitigate sequencing noise, calculating a tricube-smoothed *G’* statistic against an empirically modeled null distribution. Within *QTLseqr*, we controlled for multiple testing by estimating the False Discovery Rate (FDR) to establish a significance threshold of q-value <= 0.05.

### Functional Annotation

To predict the functional consequences of the variants identified in our BSA dataset, we used *SnpEff*^21^. We first constructed a custom database for *A. distichus* using the reference genome assembly^22^ and its corresponding GFF3 gene models. This species-specific database was then used to annotate the filtered VCF file from the BSA dataset. Each variant was classified based on its genomic location (e.g., exonic, intronic, intergenic), and its predicted functional impact on protein-coding genes was categorized as high (e.g., stop-gain), moderate (e.g., non-synonymous substitution), low (e.g., synonymous substitution), or modifier (e.g., downstream variant). This allowed us to identify specific non-synonymous variants within candidate genes for further analysis.

### Copy Number Variant Identification

To detect structural variations segregating between the dewlap morphs, we analyzed the BSA dataset utilizing *CNVpytor*^23^. To accommodate the complex neo-sex chromosome architecture and prevent coverage biases, read depth signals were processed independently for the autosomes, the X-linked scaffolds, and the neo-Y-linked scaffolds. Read depth signals were extracted across 1 kbp, 10 kbp, and 100 kbp bins. To accurately detect variants against the species’ background, independent baseline coverages were computed for the autosomes, the X-linked scaffolds, and the neo-Y-linked complex using the interquartile mean of core read depth, excluding outlier spikes. Putative CNVs were called independently for the orange and yellow pools. Finally, we isolated functionally divergent CNVs by cross-referencing the pool-specific calls utilizing *bedtools*^24^. Structural variants were classified as divergent if they exhibited strictly pool-specific occurrence or demonstrated a >= 25% divergence in physical size or normalized read depth between the morphs.

### Neo-Y-Chromosome Gene Annotation and Classification

To characterize the coding potential and structural integrity of the non-recombining Y chromosome, we identified neo-Y-linked genes and X-Y gametolog pairs using a dual comparative approach. First, we translated coding sequences from the *A. distichus* reference genome and compared them to the annotated genomes of outgroup species (*Anolis carolinensis* and *Anolis sagrei*) using *OrthoFinder*^25^ to establish orthologous gene groups. The physical coordinates of these orthogroups were then mapped across species to evaluate conserved synteny using *GENESPACE*^26^. Second, to explicitly identify X-Y gametolog pairs within *A. distichus*, we extracted all translated peptides mapping to the defined X-linked (scaffolds 8 and 12X) and neo-Y-linked (scaffolds 12Y, 13 and 16) regions. We performed mutual reciprocal best-hit *BLASTP* searches between these isolated X and Y peptide databases (e-value <= 1e-10).

By integrating the outgroup synteny data with the internal *BLASTP* alignments, we classified the *A. distichus* neo-Y-linked genes into distinct evolutionary categories: (i) X-Y gametologs, (ii) neo-Y-restricted genes lacking an X-linked counterpart, and (iii) multi-chromosomal amplicons (putative transposable elements). Furthermore, we assessed the degradation status of each neo-Y-linked gene by comparing its peptide length to the median length of its outgroup orthologs; genes measuring less than 80% of the ancestral outgroup length were classified as putative pseudogenes. Finally, to facilitate downstream analysis, the NCBI gene symbols derived from the *A. carolinensis* and *A. sagrei* outgroups were transferred to the homologous *A. distichus* loci.

### Candidate Variant Identification and Annotation

To generate a high-confidence list of candidate variants, we built an integrative computational pipeline using *bedtools*^24^ and custom *awk*, and *python* scripts to cross-reference our genomic, spatial, and transcriptomic datasets. We retained loci as candidates if they met one of two statistical thresholds: (i) they showed a significant difference in allele frequencies between the orange and yellow pools in the BSA dataset (*QTLseqr*; *q* < 0.05) and were located within 25 kbp of both a GWA locus (*p* < 0.05) and a clinal locus (AF >= 0.3); or (ii) they emerged as extreme outliers in the variant-by-variant PoolSeq analysis (*QTLseqR*; top 0.01%). We annotated the functional impact of variants within these consensus regions by crossing them with the annotated vcf file produced by *SnpEff*^21^ (see above). To isolate biologically relevant targets, we extracted divergent alleles across the cline (DeltaAF >= 0.3) and cross-referenced the annotated genes against two *a priori* lists: a dataset of differentially expressed genes between orange and yellow morphs^27^, and a curated database of known vertebrate coloration genes. Finally, we mapped the divergent structural variants identified in our CNV analysis onto these candidate regions to evaluate potential *cis*-regulatory disruptions.

### Genetic Architecture and Epistasis Modeling

To evaluate the quantitative genetic architecture of the dewlap polymorphism and test for epistasis, we modeled the statistical interaction between the neo-Y-linked haplotype and the autosomal candidate regions. We extracted all variant genotypes within a 50 kbp window (+– 25 kbp) centered on the *CYP2J2* and *TMEM68* gene clusters using *BCFtools*^10^. Missing genotypes were mean-imputed, and invariant sites were removed. To collapse the high-dimensional genotypic data of each candidate window into a single continuous predictor, we performed a Principal Component Analysis (PCA) and extracted the first principal component (Local PC1). The overall neo-Y-chromosome haplotype was similarly represented by the first principal component of all neo-Y-linked variants.

We tested for epistatic interactions by fitting independent linear models for each candidate region, evaluating dewlap hue against the interaction term between the Local PC1 and the neo-Y-haplotype PC1. Statistical significance and variance partitioning were assessed using an analysis of variance (ANOVA) using custom scripts in *R*. Following the statistical rejection of the epistatic interaction terms, we constructed a joint additive linear model incorporating the Local PC1 of *CYP2J2*, the Local PC1 of *TMEM68*, and the neo-Y-haplotype PC1. We determined the best fit model through a likelihood ratio test (LRT) of nested models with different combinations of predictors (neo-Y-haplotype, *TMEM68*, *CYP2J2*). The proportion of total phenotypic variance explained by each independent genomic component was calculated using the sequential sum of squares derived from the joint *ANOVA* table.

## Acknowledgements

Research reported in this publication was made possible in part by the services of the KU Genome Sequencing Core This lab is supported by the National Institute of General Medical Sciences (NIGMS) of the National Institutes of Health under award number P20GM103638. We thank the Ministerio de Medio Ambiente y Recursos Naturales and the Museo Nacional de História Natural of the Dominican Republic for collecting and exportation permits and logistic help. We thank Cristian Marte, Eveling Gabot, Patricia Pineda, Javier Torres and Tanner Myers for support in field work.

## Author Contributions

P.L.H.d.M. and R.E.G. conceptualized the project and collected the specimens. P.L.H.d.M. collected the molecular data and performed the statistical analyses, with analytical design input from J.K.K. P.L.H.d.M. wrote the initial draft of the manuscript, and all authors contributed to the writing and editing of the final version.

## Materials & Correspondence

### Data availability

The data supporting the findings of this study are available within the Article itself and its Supplementary Information. Sequencing reads (WGS, MSG, and BSA) are deposited in the NCBI Sequence Read Archive under BioProject [TBD]. Phenotypic data (images, spectrophotometry, histology) and filtered VCF files are deposited in Dryad under DOI [TBD].

## Code Availability

Custom scripts used for image processing, VCF filtering, read-depth normalization, and epistatic modeling are available on GitHub at [TBD] and archived on Zenodo under DOI [TBD].

## Library Preparation and Sequencing

### DNA Extraction

We extracted DNA from tissues with *Speedbeads*^1^ following the protocol available at http://phyletica.org/lab-protocols/extraction-spri.html. Briefly, we digested approximately 0.1µg per sample in 200µl of lysis buffer with 5% proteinase K (Thermo Fisher) in a 2ml microcentrifuge tube for at least 12 hrs. The extraction protocol consists of four main steps: binding DNA to beads, separating DNA in beads to contaminants, washing beads and DNA with cold 70% ethanol to remove contaminants, and eluting DNA fragments from beads.

We bound DNA to *Speedbeads* by combining equal volumes of Speedbeads and digestion product (1:1, v/v), flicking the mixture, and incubating it in the dark for 5 minutes. Samples were then placed in a magnetic stand (Fisher Scientific) for 5 minutes in the dark. The beads were washed twice with 400µl of cold (4°C) 70% Ethanol. Each wash involved a 30-second incubation before aspirating and discarding the Ethanol. Samples were subsequently dried in a heat block at 37°C for 5 minutes. For re-elution, we added 200µl of EB (Qiagen), flick-mixed and incubated in the dark for 5 minutes. Samples were again placed in a magnetic stand for 5 minutes in the dark, and the resulting eluate was transferred to new, individually labeled 2ml microcentrifuge tubes. Quality control of the samples was performed by quantifying DNA concentration using a QubitHS (Invitrogen) and assessing purity with NanoDrop (Thermo Scientific).

### Library Preparation

We acquired genomic data for our genome-wide association (GWA) and geographic cline analyses through a reduced representation sequencing library preparation for multiplexed shotgun genotyping (MSG) described in Andolfatto et al.^2^. Our objective was to sample approximately 10% of each sample’s genome at 10x coverage. Given studies using genome-wide markers^3^ we anticipated linkage disequilibrium to be low for specimens across the transect. To achieve this, we utilized a custom *python* 2.7 script to *in silico* digest the Hispaniolan Bark Anole genome^4^ with various commercially available restriction enzymes. Based on this analysis, we selected BfaI (New England Biolabs) as the optimal restriction enzyme for our study.

We performed the MSG library preparation at the KU Genome Sequencing Core (GSC). Library preparation was divided into six steps: digestion, barcode ligation, post-ligation bead cleaning, size selection, post-size selection bead cleaning, PCR amplification, and PCR purification and quality control. We first normalized genomic DNA samples to 2 ng/µl using EpureH2O. For digestion, we added 5 µl (10 ng) of each sample to a solution containing 2 µl of NEB buffer 4, 0.2 µl of BSA (100x), 0.3 µl of BfaI (NEB), and 12.5 µl of EpureH2O. Digestion proceeded for 3 hours at 37°C, followed by enzyme inactivation for 20 minutes at 65°C. We ligated barcoded adapters by adding 1 µl of adapter oligonucleotides (5 µM), 5 µl of 10x T4 DNA ligase buffer (NEB), 0.38 µl of T4 DNA Ligase (150U, NEB), and 23.62 µl of EpureH2O to each sample. Ligation occurred in a thermocycler (Bio-Rad) at 25°C for 2 hours, followed by enzyme inactivation for 10 minutes at 65°C. We then pooled and precipitated ligation products using 3M sodium acetate (pH 5.2) and isopropanol (1:10, v/v). We resuspended the dried pellet in 100 µl of 1x TE buffer (Invitrogen) by heating it at 65°C for 30 minutes. We extracted suspended DNA with phenol:chloroform (1:1, v/v) and purified it using 1.5 volumes of AMPure XP beads (Beckman Coulter). After 5 minutes incubation at room temperature, beads were captured on a magnet for 10 minutes, the supernatant discarded, and the beads washed twice with 70% ethanol. Beads were dried for 4 minutes at 37°C before eluting DNA in 32 µl of 1x TE buffer.

We size-selected DNA pools for a 250-300 bp range using a Pippin Prep instrument (Sage Science) with a 2% agarose cassette and internal standards. Following size selection, we exchanged buffers for 2 volumes of AMPure XP beads per 1 volume of sample, following the standard AMPure protocol. We quantified size-selected DNA using a Qubit HS assay (Invitrogen). We PCR amplified 2 ng of the size-selected DNA in a 50 µl reaction containing 10 µl of 5x Phusion Buffer (NEB), 1 µl of 10 mM dNTPs, 2.5 µl of FC1 primer (10 µM), 2.5 µl of FC2 primer (10 µM), 0.5 µl of Phusion High-Fidelity DNA Polymerase (NEB), and 33.5 µl of EpureH2O. We purified PCR products using the Agencourt AMPure PCR purification kit (Agencourt) with a 0.8x bead concentration and eluted in 30 µl of 10 mM Tris buffer (NEB). We used 2 µl of the purified product for final quantification with a Qubit HS assay. We then stored resulting libraries at –80°C before shipping to Novogene for sequencing on an Illumina HiSeqX platform (150 bp paired-end reads across three lanes).

For the bulk segregant analysis (BSA) we picked 50 specimens from either extreme of the hue spectrum to assemble pools. We independently equilibrated three technical replicates per pool to reduce the effects of amplification biases^5^. Each pool consisted of 1 µg of DNA equally distributed across the 50 specimens. We kept technical replicates at –80°C until ready for shipping. The whole genome shotgun (WGS) library was prepared with a Swift Ws Turbo DNA library kit protocol (Swift Biosciences) and sequenced in a Novaseq 4000. BSA samples were sequenced to approximately 16.5x coverage per replicate, and WGS samples were sequenced to approximately 20x per sample. BSA and WGS library preparation and sequencing was performed at the Oklahoma Medical Research Facility (OMRF).

## Bioinformatics

### Read Processing and Alignment

We demultiplexed and quality filtered raw reads from the MSG dataset using the process_radtags module in *STACKS* (v2.62)^6^, discarding reads with uncalled bases or low-quality scores (Phred < 20) and rescuing barcodes/RAD-tags with up to one mismatch. We processed raw reads from the WGS and PoolSeq datasets using *fastp*^7^ to remove adapters and low-quality sequences with default parameters. We aligned all processed reads to the *A. distichus* reference genome^4^ using *BWA*^8^ with default settings. We converted alignments to BAM format, sorted, and filtered for mapping quality (MAPQ ≥ 30) using *SAMtools*^9^.

### Variant Calling and Initial Subsetting

We performed joint variant calling across all samples using *BCFtools* ‘*mpileup*,’ and ‘*call*’^8^, skipping indels and retaining allele depth (AD) and total depth (DP) information. We then subsetted the resulting multi-sample BCF file using *BCFtools* ‘*view*’ to create four distinct datasets: WGS (2 males, 2 females for sex chromosome analysis), BSA (orange and yellow pools), GWA (individuals from the contact zone center), and Cline (all individuals across the transect).

### Dataset-Specific Filtering

Each dataset underwent specific filtering:

– **WGS Dataset:** Filtered using a custom *python* script to retain invariant and bi-allelic loci with mapping and call quality ≥ 20, present in at all 4 individuals.
– **BSA Dataset:** Split into two subsets. The Association subset retained only bi-allelic variant sites present in both pools with MQ ≥ 20, QUAL ≥ 20, and MAF ≥ 0.05. The Population subset used the same filters but also retained invariant sites.
– **GWA Dataset:** Filtered using a modified custom *python* script to retain bi-allelic SNPs with MQ ≥ 20, QUAL ≥ 20, MAF ≥ 0.20, and present in at least 60 of the 195 individuals.
– **Cline Dataset:** Filtered using a stepwise strategy implemented with *VCFtools*^10^. We first removed sites with >50% missing data and individuals with >90% missing data. We then removed indels, non-biallelic sites, and sites deviating from Hardy-Weinberg equilibrium (*p* < 0.01). Subsequently, we explored stricter thresholds for site and individual missingness (60%/70%, 70%/50%, 80%/30%, 90%/20%, 95%/15%).

### Imputation

As is characteristic of reduced-representation sequencing, the final GWA and Cline genotype matrices exhibited varying levels of missing data across individuals. We imputed these missing genotypes using *BEAGLE* (v.5.5;^11^) with default parameters. We assessed the sensitivity of our analyses to missing data by imputing multiple filtering iterations of the Cline dataset. As the clinal patterns remained consistent regardless of filtering stringency, the presented results focus on the dataset derived from the baseline 30% per locus and 20% individual missingness thresholds.

## Characterizing the Degradation of the Neo-Sex Chromosomes of *A. distichus*

### Rationale and Predictions

The *Anolis distichus* genome features an XXY sex chromosome system derived from multiple autosome-to-sex chromosome fusions^4^. Theoretical models demonstrate that in populations characterized by differential migration between males and females, sex chromosomes contribute disproportionately to local adaptation, accumulating locally adapted alleles at a significantly higher rate than autosomes^12^. As these diverging sex-linked sequences cease recombining and structurally degrade, they provide a hemizygous environment that further shields these co-adapted alleles from homogenizing gene flow. Given the recency of these fusions in *A. distichus*^13^, we characterized the extent of recombination suppression and structural hemizygosity across the Y and neo-Y chromosomes to identify the boundaries of this isolated genomic architecture.

We predicted specific genomic signatures based on chromosome state^14–16^:

– **Hemizygous X Regions**: Expected to show a ∼2-fold higher read depth in females vs. males (log2[F:M] ≈ 1), higher heterozygosity in females (log10[Het F:M] > 0), and a copy number relative to autosomes of ∼3/4 (3 X copies per 4 autosome copies in a 2F:2M sample).
– **Fully Degraded Y Regions:** Expected to show strongly male-biased read depth (log2[F:M] < –1), elevated linkage disequilibrium (LD) due to lack of recombination, and a copy number relative to autosomes of ∼1/4.
– **Pseudo-Autosomal Regions (PARs) / Non-degraded Y homologs:** Expected to show similar read depth between sexes (log2[F:M] ≈ 0) and similar heterozygosity (log10[Het F:M] ≈ 0). Copy number relative to autosomes depends on assembly: ∼0.5 if assembled separately on X and Y scaffolds, ∼1 if collapsed onto one scaffold.
– **Regions Undergoing Degradation:** Expected to show intermediate patterns (e.g., 0 < log2[F:M] < 1 for X-linked regions) and potentially elevated “heterozygosity” due to mismapping of divergent reads.

### Approach

We used WGS reads from two males and two females to obtain the data for our assessment of sex chromosome associated regions of the genome. We obtained per-base read depth using *SAMtools* ‘*depth*’^9^ function. To normalize across samples, we calculated the mean autosomal depth for each individual and then computed a weighted mean depth per base pair using custom *awk* scripts. We then calculated mean statistics across 10 kbp sliding windows (1 kbp steps) using *BEDTools* ‘*getwindow*’^17^ and custom *awk* scripts. Because this standard metric does not fully capture the extent of Y degeneration, we developed a weighted depth-of-coverage ratio that normalizes the combined female and male depth against the mean autosomal depth of the same samples. Theoretically, fully pseudoautosomal regions will exhibit a ratio of ∼1.0, completely hemizygous X-linked regions a ratio of ∼0.75, and strictly neo-Y-linked regions a ratio of ∼0.25. By applying this normalized metric, we could explicitly identify active regions of degradation falling between the ∼0.75 and ∼1.0 thresholds.

Following the suppression of recombination, degrading sex chromosomes are expected to accumulate private alleles and fixed X-Y differences, the latter of which manifest as elevated male heterozygosity due to the cross-mapping of divergent Y reads^15^. To capture these genomic signatures of degradation, we calculated the proportion of heterozygous sites, total segregating SNPs, and sex-specific SNPs per window for each individual. To explicitly evaluate sex-biased heterozygosity and SNP density, we calculated a normalized difference index defined as (mean(Xf) – mean(Xm))/max(mean(Xf), mean(Xm)), where mean(Xf) and mean(Xm) represent the mean window values for females and males, respectively. For the density of sex-specific SNPs, we calculated the unnormalized difference between the female and male proportions. All statistical transformations and windowed visualizations were performed using custom scripts in *R*.

To corroborate depth patterns, we repeated the Sex:Autosome depth comparison using the male-only GWA dataset (MSG reads). Finally, to confirm the non-recombining nature of putative Y regions, we calculated linkage disequilibrium (LD) using the GWA dataset and *PLINK* (v1.9)^18^ with flags “--r2 gz –-ld-window-kb 1000000 –-ld-window-r2 0.” Mean LD decay and LD heatmaps were generated per chromosome using custom R scripts.

### Identification and Partitioning of the Chimeric Scaffold 12

Initial whole-genome sliding-window analyses of read depth and genetic variation revealed a sharp structural transition within scaffold 12, suggesting a chimeric assembly of distinct sex-linked regions. To map the evolutionary boundary of this scaffold, we profiled the log2-transformed female-to-male read depth ratio, the normalized female-to-male heterozygosity index, the density of sex-specific SNPs, and the Sex:Autosome depth ratio (see above). We identified an abrupt genomic transition at the 8.7 Mbp locus. Across this boundary, the chromosomal signature shifts definitively from a highly degraded, neo-Y-linked profile (characterized by male-biased read depth, a high density of male-specific SNPs, and a Sex:Autosome ratio ∼ 0.25) to a hemizygous X-linked profile (characterized by a 2:1 female-to-male depth ratio, female-biased heterozygosity, and a Sex:Autosome ratio ∼ 0.75; Supplementary Fig. 1).

To contextualize this transition, we looked for assembly “scars”—defined as continuous runs of 100 or more undetermined nucleotides (’N’)—across the reference genome to identify putative scaffolding junctions. We found an exact alignment of the 8.7 Mbp genetic transition with a major scaffolding scar providing initial evidence of an artificial fusion. Next, we corroborated this physical boundary using Hi-C chromatin interaction maps. Balanced contact frequencies were extracted at a 25 kbp resolution, log2p-transformed, and mapped across a 4 Mbp genomic window centered on the putative breakpoint. A distinct discontinuity in chromatin contact frequencies exactly at the 8.7 Mbp scaffolding scar provided orthogonal validation of the assembly error. Lastly, self-alignment of the partitioned scaffold segments using NUCmer (*MUMmer* v4.0) confirmed distinct internal synteny profiles. Based on this concordant physical read depth, sequence variance, scaffolding topography, chromatin conformation, and synteny, we established 8.7 Mbp as the definitive evolutionary breakpoint. Consequently, was programmatically partitioned into independent *12Y* (< 8.7 Mbp) and *12X* (> 8.7 Mbp) domains prior to all downstream population genomic and quantitative association analyses.

### Histological Sample Preparation and Imaging

Histological analyses were performed on four male *A. distichus* from a captive colony at the University of Kansas (KU), following protocols approved by the KU IACUC (AUS 208-03). All electron microscopy (EM) processing and imaging were conducted at the KU Microscopy and Analytical Imaging (MAI) laboratory (RRID:SCR_021801).

### Scanning Electron Microscopy (SEM)

For SEM, 1×1 cm skin samples were dissected from the dewlap (orange and yellow), belly (white), and back (green) of one specimen. Samples were fixed overnight at 4°C in 4% paraformaldehyde (PFA) in 1X Hanks’ Balanced Salt Solution (HBSS). The following day, samples were post-fixed in modified Karnovsky’s fixative (2.5% glutaraldehyde / 2% PFA in 0.2M sodium cacodylate buffer, pH 7.4) for 1 hour, rinsed twice (5 min each) in 0.2M sodium cacodylate buffer, and stained with 0.5% Osmium tetroxide in 0.2M cacodylate buffer for 2 hours at room temperature. Due to the hydrophobic nature of the skin, fixation and staining were performed using reagent vapors. Samples were then rinsed twice (5 min each) in cacodylate buffer, transferred to sample holders, immersed in methanol, and critical point dried using a K850 Critical Point Drier (Quorum Technologies).

Dried SEM samples were mounted on aluminum stubs with carbon tape. Images were acquired on a Hitachi SU8230 Cold Field Emission SEM using a YAG backscattered electron detector at 5.0 kV accelerating voltage and 8.0 mm working distance. High-resolution images (2560 x 1920 pixels) were captured with 32-64 s scan speeds. Lower magnification overview images were also taken using secondary electron detectors.

### Transmission Electron Microscopy (TEM)

For TEM, 1×1 cm skin samples were dissected from the dewlap, belly, and back of the three remaining specimens. For individuals with bicolored dewlaps, the orange center and yellow margin were processed separately. Samples were fixed overnight at 4°C in 4% PFA in 1X HBSS. The next day, tissues were subsampled into 2×2 mm squares and post-fixed in a solution containing 2.5% glutaraldehyde, 2% PFA, and 0.5% Osmium tetroxide in 0.2M cacodylate buffer (pH 7.4) for 30 min at room temperature. Osmium staining was enhanced by rinsing in 50% ethanol (5 min) followed by treatment with 1% p-phenylenediamine in 70% ethanol (5 min). Samples were then dehydrated through an ethanol series (95% 5 min; 100% 2× 5 min) followed by 1:1 ethanol:acetone (5 min). Dehydrated samples were infiltrated with Embed 812 resin through a series of acetone:resin mixtures (1:1 5 min; 1:3 5 min) followed by two changes of 100% Embed 812 resin (15 min each). Samples were immersed in Embed 812 resin + DMP-30 catalyst in flat molds and polymerized at 65°C for 48-72 hours.

For initial survey, thick sections (500 nm) were cut using a diamond knife on a Sorvall JB-4A microtome, stained with 25% Toluidine Blue on a hot plate (200°C, 3-5 s), and observed under a Leica DM750 light microscope. Subsequently, thin sections (70-90 nm) were cut using a Leica UC6 ultramicrotome, mounted on G300-Cu copper grids, and stained sequentially with Uranyl acetate (UranyLess, 15 min) and 3% Lead Citrate (5 min in a CO2-depleted atmosphere), with appropriate water rinses between stains. TEM images were acquired on a Hitachi H-8100 TEM at 200 kV using a BIOSPR16 camera, with magnifications ranging from 1500x to 4000x.

### Reagents

Paraformaldehyde (PFA; Sigma Aldrich, St. Louis, MO, Cat. No. P6148, Lot. No. MKCD5277); sucrose (Fisher Chemical, Waltham, MA, Cat. No. 57-50-1, Lot. No. 166358); Hanks’ Balanced Salt Solution (HBSS; Sigma Aldrich, St. Louis, MO, Cat. No. 14185052, Lot. No. 2042301); Glutaraldehyde (25% in aqueous solution; Electron Microscopy Sciences [EMS], Ft. Washington, Cat. No. 16216, Lot. No. 2170117); Sodium Cacodylate Trihydrate (EMS, Cat. No. 12300, Lot. No. 980908); Osmium tetroxide (EMS, Cat. No. 19112, Lot. No. 170711-03); p-phenylenediamine (Sigma Aldrich, Cat. No. P6001/ Lot. No. WXBB8077V); Uranyl acetate (UA; EMS, Cat. No. 22400, Lot. No. 100606); Lead citrate (LC; EMS, Cat. No. 17800, Lot. No. 060630); Phosphotungstic acid (PTA; EMS, Cat. No. N/A/. Lot. No. N/A); Embed 812 resin (EMS; Embed 812 resin consist of: (1) Embed 812, Cat. No. 14900, Lot. No. 921120; (2) Dodecenyl Succinic Anhydride, Cat. No. 13700, Lot. No. 970325; (3) Nadic Methyl Anhydride, Cat. No. 19000, Lot. No. 970325); the catalyst for embedding with Embed 812 resin: DMP-30 (EMS, Cat. No. 13600, Lot. No. 90-72-2).

## Pigment Biochemistry

### Background

Yellow, orange, and red coloration in vertebrates is primarily driven by two distinct classes of pigments: carotenoids and pteridines^26,27^. Because vertebrates lack the enzymatic pathways to synthesize carotenoids *de novo*, these pigments must be acquired from the diet; in contrast, pteridines are synthesized endogenously^28,29^. In *Anolis* lizards, the transition from yellow to orange and red skin is typically mediated by increased concentrations of dietary carotenoids (e.g., lutein and zeaxanthin), the localized deposition of red-absorbing pteridines (e.g., drosopterins), or a combination of both^30,31^. To determine the chemical basis of the dewlap polymorphism in *A. distichus*, we sequentially extracted these two pigment classes from skin samples and quantified their relative abundance using absorbance spectroscopy.

### Skin Patch Sampling and Pigment Extraction

We sequentially extracted carotenoids and pteridines from back, belly, and dewlap skin samples using a modified version of the protocol by McLean et al.^32^. All procedures were performed under dim red light. Prior to extraction, 1×1 mm skin samples were weighed using a high-precision scale (Mettler Toledo).

For carotenoid extraction, samples were placed in 2 ml screw-cap tubes with 2.2 mm Tungsten Carbide Milling Media balls (MSE Supplies) and 500 µl of methanol:ethyl acetate (6:4, v/v) containing 0.01% butylated hydroxytoluene (BHT) to prevent oxidation. Samples were homogenized twice in a Qiagen TissueLyser (30 Hz, 1.5 min each), incubated on ice in a shaker (350 rpm) for 30 min, and centrifuged (Thermo Scientific ST 16R, 15200 rpm, 4°C). The supernatant was transferred to a new foil-covered tube and stored temporarily at –20°C. The pellet was re-extracted with 200 µl of the same solvent, following the same incubation and centrifugation steps. The supernatants from both extractions were pooled, the tube flushed with argon, flash-frozen in liquid nitrogen, and stored at –80°C.

Pteridines were subsequently extracted from the remaining pellet by adding 500 µl of 2% ammonium hydroxide. The homogenization, incubation, and centrifugation steps were repeated as for carotenoids. The pellet was re-extracted with 200 µl of 2% ammonium hydroxide. Both pteridine supernatants were pooled, flushed with argon, flash-frozen, and stored at –80°C. Blank extractions (solvent only) were performed alongside samples for spectrophotometer standardization.

### Absorbance Spectroscopy

For each extract (carotenoid and pteridine), 200 µl was transferred to a quartz cuvette. Absorbance spectra (UV-VIS-NIR) were measured using an absorbance spectrophotometer relative to the corresponding blank solvent. To ensure linearity and reduce noise, samples with absorbance > 2 arbitrary units (AU) were serially diluted tenfold until absorbance fell below this threshold. Absorbance data were plotted using custom *R* scripts utilizing the ‘ggplot2’ and ‘tidyverse’ packages^33^.

### Computational Pipeline and Statistical Software

All data processing, visualization, and statistical analyses were conducted in R version 4.3.0 or higher. The analytical workflow was primarily constructed using the *tidyverse* suite of packages^33^, with *readr* and *dplyr* utilized for data ingestion and manipulation, and *ggplot2* for high-resolution graphics. Automated file system traversals across nested instrument output directories were managed via the *fs* package.

### Spectral Data Acquisition and Pre-processing

Skin pigment extracts were analyzed using visible light spectrophotometry (400–700 nm). To ensure analytical integrity and prevent instrument-driven bias, a multi-step pre-processing pipeline was implemented. Raw spectra were first baseline-corrected by subtracting the minimum absorbance value within the 400–700 nm range to ground each curve at zero (A_cor = A – min(A)). A critical challenge in pigment spectrophotometry is detector saturation at high concentrations (typically >1.5 AU), which results in “flat-top” peaks that artificially distort the spectral shape and underestimate total absorbance. To mitigate this, for every specimen-tissue combination where multiple scans existed, we prioritized the file with the highest dilution factor (e.g., 2X-Diluted over Standard). This ensured that all spectral shape analyses were performed on non-saturated curves where the primary absorbance peaks remained within the linear dynamic range of the detector.

### Spectral Shape Analysis (Composition)

To isolate chemical composition from concentration, baseline-corrected spectra were maximum-normalized (A_norm = A_corr / max(A_corr)). Shape variance was evaluated using a centered Principal Component Analysis (PCA) on the normalized spectral matrix. To determine if pigment composition differed across phenotypes, specimens were categorized into four comparison groups: Back skin, Belly skin, Yellow Dewlap, and Orange Dewlap. The primary axis of variation (PC1) served as the metric for compositional difference. The statistical significance of these groupings was tested using a one-way ANOVA followed by Tukey’s Honest Significant Difference (HSD) post-hoc test.

### Reconstruction of True Absorbance (Concentration)

Total pigment abundance was quantified using the Area Under the Curve (AUC), calculated via trapezoidal integration of the non-normalized, baseline-corrected spectra. To reconstruct the biological concentration of the original tissue from diluted samples, we applied the Beer-Lambert Law: AUC = AUC_Measured * Dilution Factor. To maintain maximum precision and minimize the propagation of error from high-fold dilutions, concentration comparisons were restricted to samples with a dilution factor <= 2. Differences in total concentration were statistically evaluated using a Welch’s two-sample t-test.

---

**Supplementary Figure 1.**
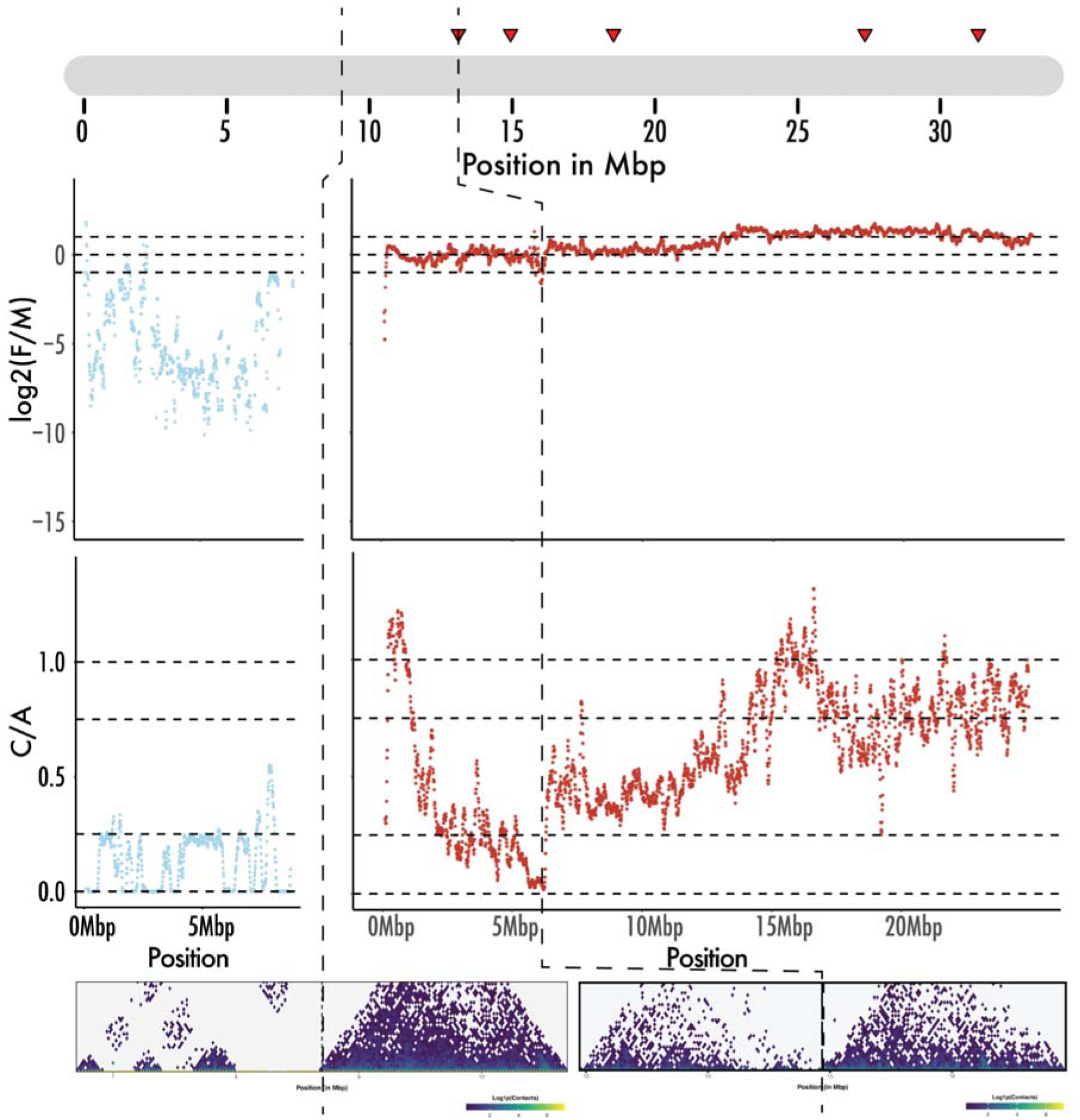
Integrative coverage and Hi-C analyses resolve the boundary between the highly degraded Neo-Y-linked and pseudoautosomal regions of Scaffold 12. (Top) Distribution of scaffolding “scars” (100 bp gaps of “N”s merged via Hi-C data). (Upper Middle) log2([F:M]) read depth from 0 to 8.7 Mbp (scaffold 12Y) and from 8.7 Mbp to the scaffold terminus (scaffold 12X). An abrupt shift at 8.7 Mbp marks the transition from strictly male-biased coverage to approximately equal, autosomal-like coverage. (Lower Middle) Normalized log2([C:A]) depth of coverage across scaffolds 12Y and 12X. (Bottom) Hi-C heatmaps for 4 Mbp windows centered at the 8.7 Mbp transition zone and the 14.94 Mbp scaffolding scar. Cross-referencing depth patterns with Hi-C data confirms the 0–8.7 Mbp region (12Y) is strictly male and highly degraded. Beyond 8.7 Mbp, the region oscillates between autosomal and male-biased coverage, and the log2([C:A]) ratio fluctuates between 0.25 and 0.5. We interpret this 12X segment as a pseudoautosomal region undergoing active degradation, and provisionally assigned it to the X chromosome assembly.

**Supplementary Figure 2.**
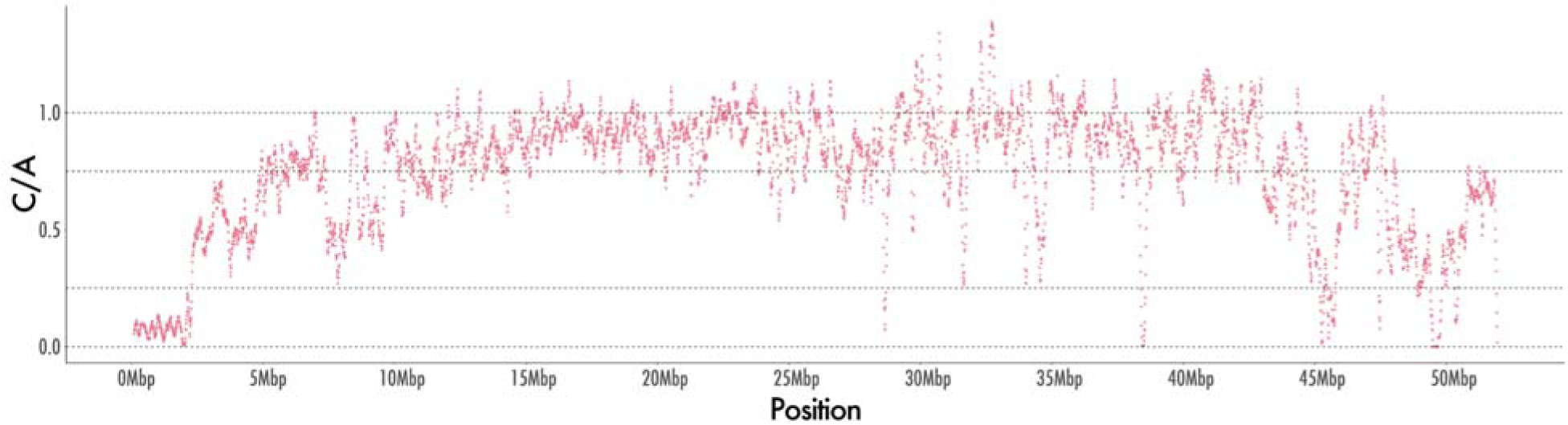
Read depth across Scaffold 8 is consistent with X-linkage. The overall low mapping of male and female reads at the beginning of the scaffold is followed by an abrupt, and then gradual, increase in relative depth towards the ∼ 0.75 ratio expected for the X. As with autosomes, the depth drops once more at the far right end of the scaffold, consistent with the lower quality of read mapping at telomeric or centromeric regions of chromosomes.

**Supplementary Figure 3.**
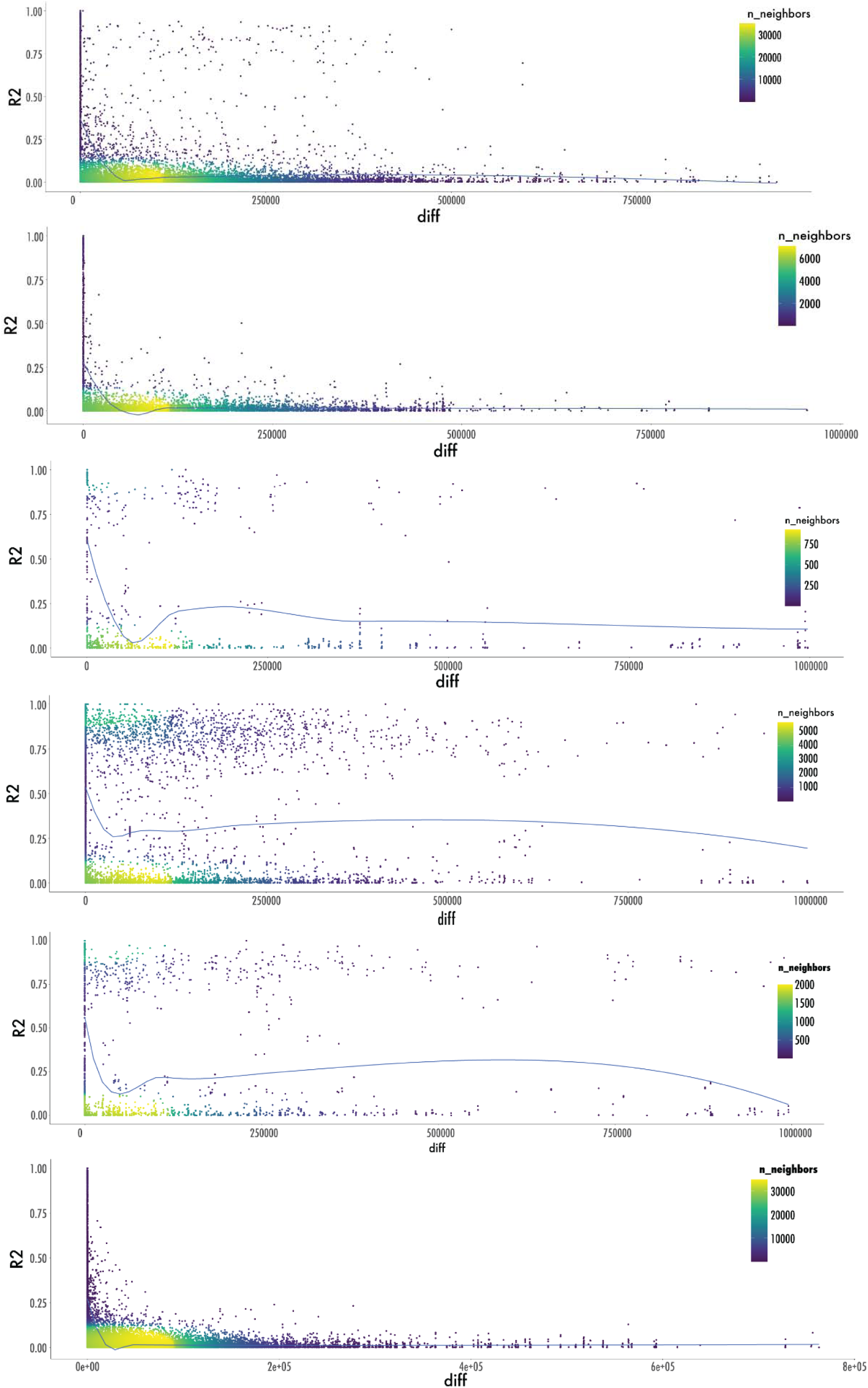
Neo-Y-linked scaffolds maintain extensive long-distance linkage disequilibrium relative to autosomes and X-linked regions. (A–F) Linkage disequilibrium (LD) decay profiles for scaffolds 8, 12X, 12Y, 13, 16, and the autosomal reference, scaffold 9. Each data point represents the squared allele frequency correlation (r2) between two GWA loci. Point density is mapped to a viridis color scale, with LOESS regression lines indicating the overarching spatial trend of r2 across physical distance. Autosomal LD (scaffold 9) decays rapidly, followed by intermediate decay rates on the X-linked scaffolds (8 and 12X). In contrast, extensive long-distance LD is maintained across the neo-Y-linked scaffolds (12Y, 13, and 16). Notably, this pronounced neo-Y-linked LD remains robustly detectable despite the prevalence of transposable elements and copy number variations, which inherently elevate genotyping noise on the Y chromosome even under strict filtering parameters.

**Supplementary Figure 4.**
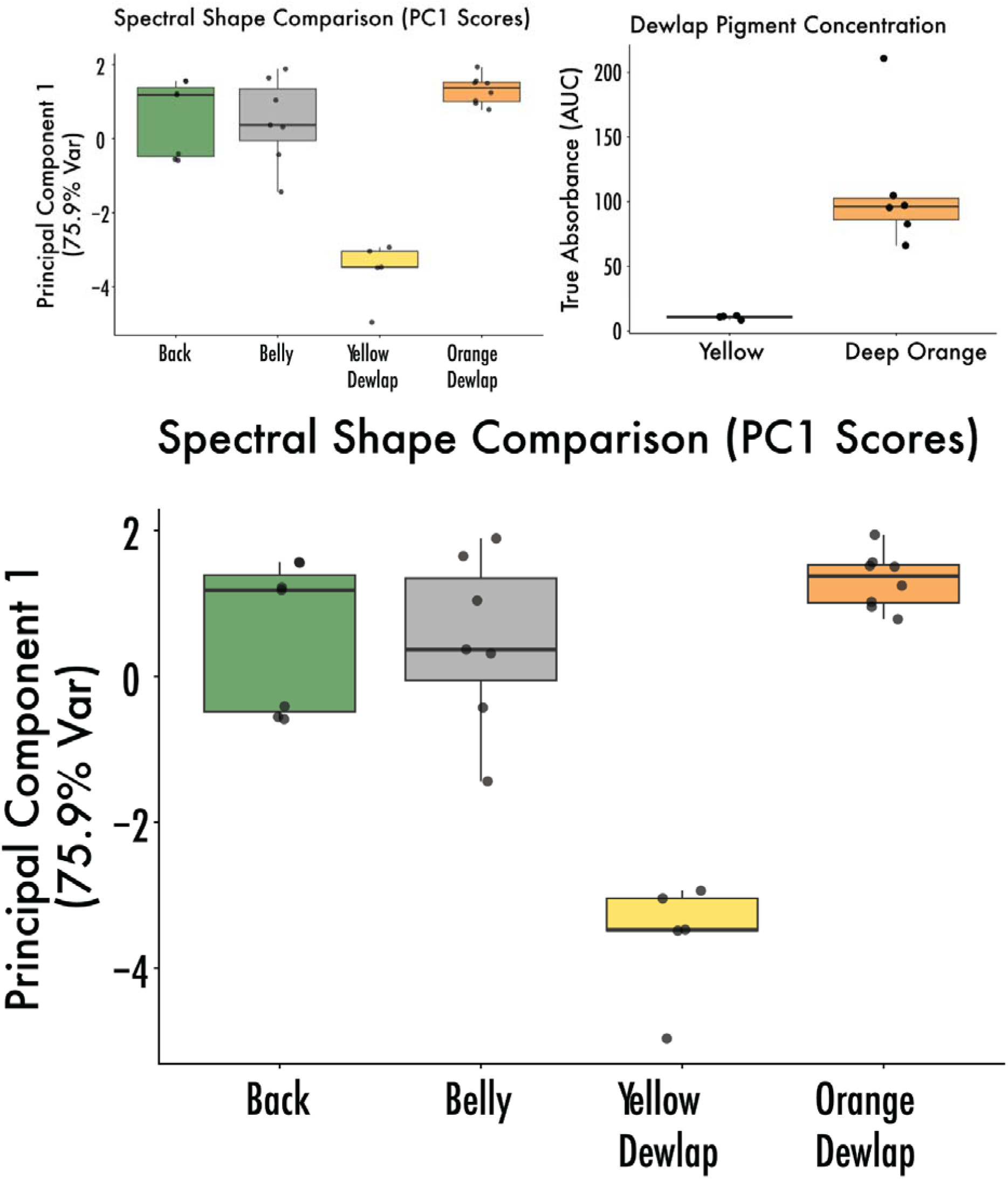
Compositional and quantitative divergence of pigments across *A. distichus* tissues. (Top Left) Boxplots of Principal Component 1 (PC1) scores derived from maximum-normalized absorbance spectra (400–700 nm). Orange dewlaps exhibit a pigment composition indistinguishable from body tissues (back and belly), whereas yellow dewlaps are compositionally distinct. **(Top Right)** Total pigment concentration estimated via the Area Under the Curve (AUC) of baseline-corrected spectra, adjusted for dilution. Orange dewlaps contain significantly higher absolute carotenoid concentrations than yellow dewlaps. **(Bottom)** Mean normalized absorbance spectra. Averaged spectral curves for the back (forest green), belly (gray), yellow dewlap (gold), and orange dewlap (dark orange) highlight a ∼13 nm rightward shift unique to the yellow phenotype.

**Supplementary Figure 5.**
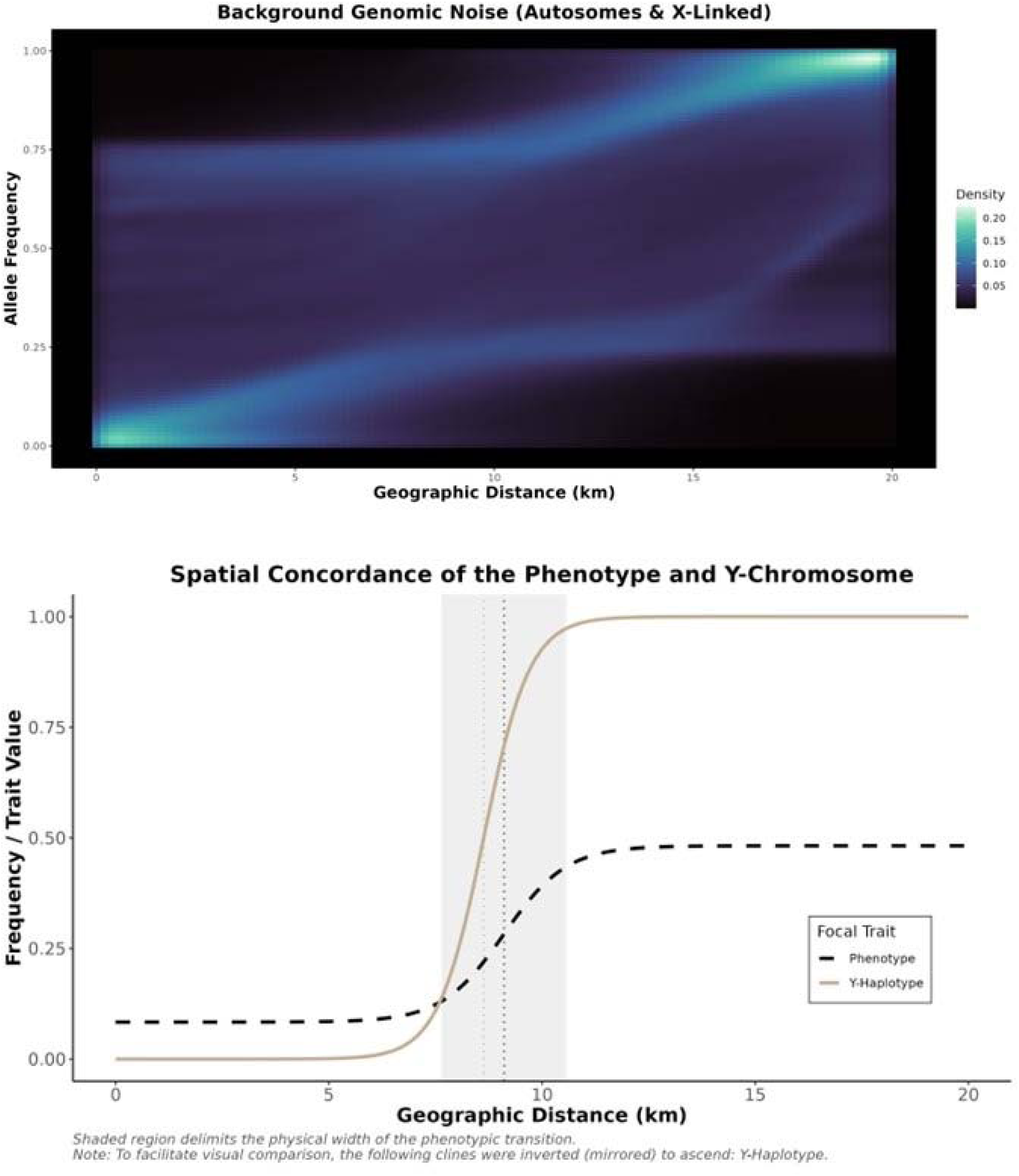
Stark spatial concordance between the neo-Y-haplotype and dewlap color demonstrates strong neo-Y-linkage despite widespread gene flow. (A) Allele frequency heatmap representing background genomic noise from autosomal and X-linked loci across the transect, with color gradients indicating point density. (B) Geographic clines representing the spatial transition of the focal phenotype and neo-Y-haplotype frequency across the sampled transect. To facilitate direct visual comparison of the cline shapes, the neo-Y-haplotype cline was inverted (mirrored) to ascend. The shaded vertical region demarcates the physical width of the phenotypic transition zone. The dashed vertical lines represent the center of the phenotypic (black) and neo-Y-haplotype (yellow) clines. The stark contrast between the sharp, overlapping neo-Y-linked/phenotypic clines and the unstructured genomic background demonstrates tight physical linkage of the phenotype to the Y chromosome, despite widespread gene flow in the rest of the genome.

**Supplementary Figure 6.**
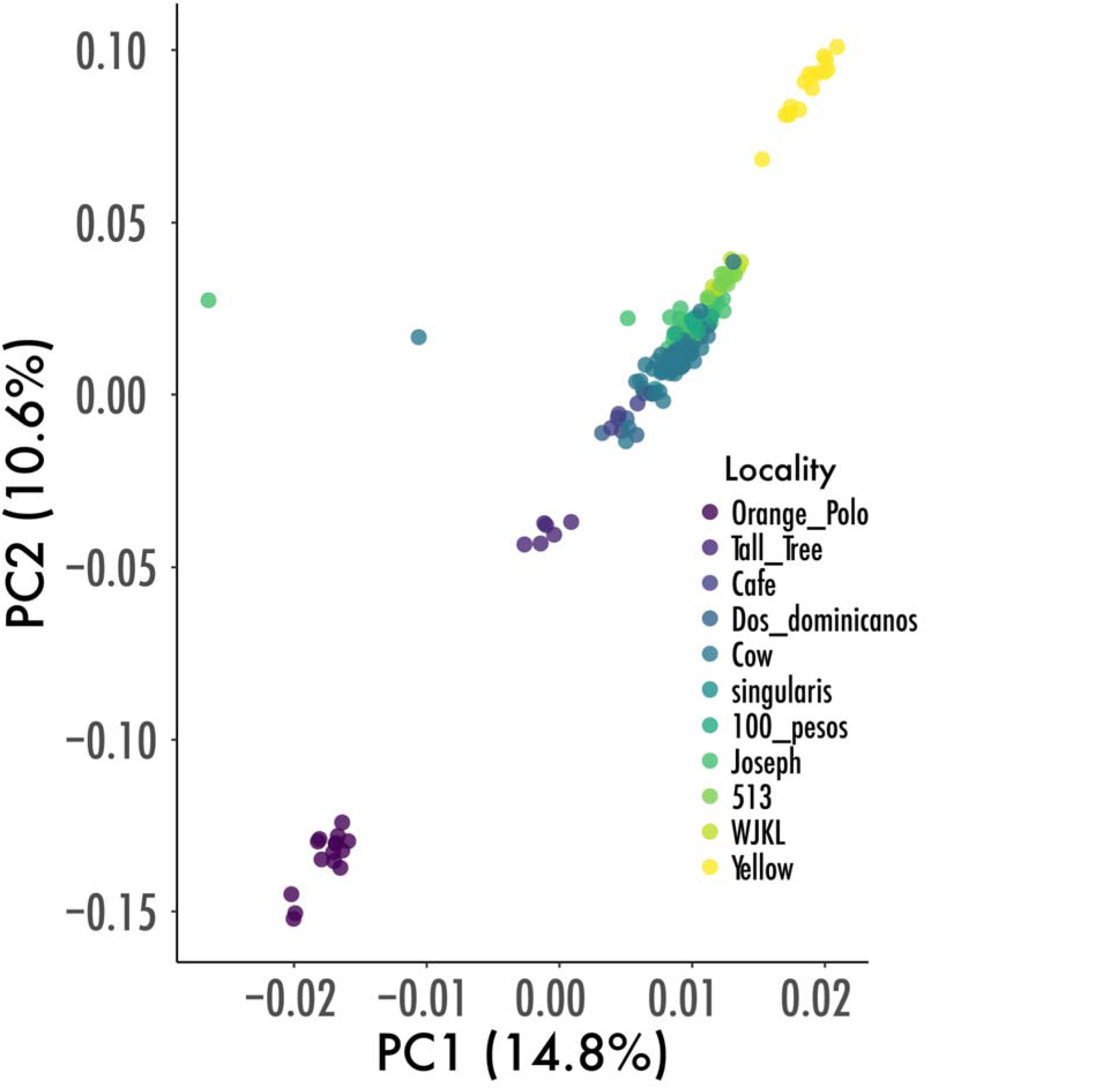
Cline Center Shows Reduced Genomics Structure. Specimens from the “contact zone” used in the GWA analyses were sampled at “Cow,” “singularis,” “100 pesos,” and “Joseph.” Orange Polo is the western-most locality (locality 1 in Fig. 1), while Yellow is the eastern-most locality (locality 11 in Fig. 1). Locality legend is ordered following number ids from Figure 1.

**Supplementary Figure 7.**
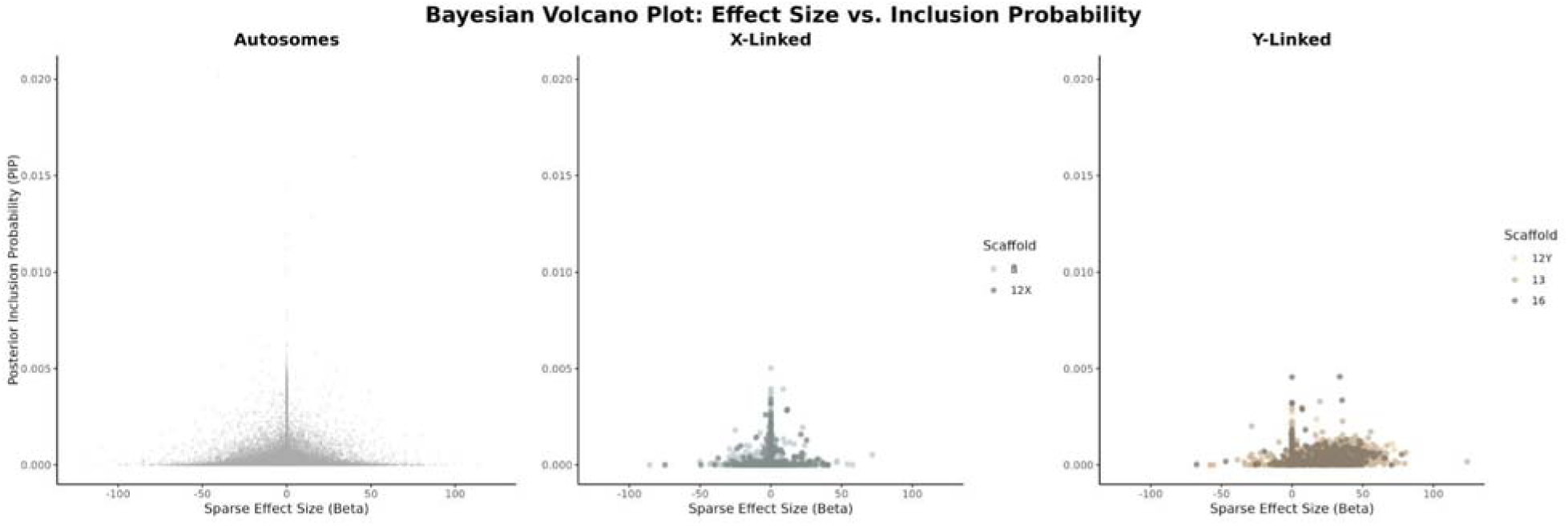
Bayesian sparse linear mixed modeling identifies the neo-Y-chromosome as the primary determinant of dewlap color pattern. Bayesian volcano plots illustrate the relationship between the sparse effect size (β, x-axis) and the posterior inclusion probability (PIP, y-axis) for individual genetic variants. Variants are partitioned by genomic linkage: Autosomes, X-linked (scaffolds 8 and 12X), and neo-Y-linked (scaffolds 12Y, 13, and 16). While autosomal and X-linked variants exhibit negligible effect sizes and baseline inclusion probabilities, the neo-Y-linked scaffolds display a distinct, consistent clustering of larger effect sizes and PIPs. The relatively low absolute PIP values observed among these top neo-Y-linked variants are an expected feature of Bayesian variable selection in regions of extensive linkage disequilibrium (see Supplementary Figure 3); because the model partitions the phenotypic variance across multiple highly correlated loci, the inclusion probability is distributed across the haplotype rather than localized to a single causal variant. This collective elevation of neo-Y-linked variants confirms a distinct, sex-linked genetic architecture underlying the color polymorphism.

## Supplementary Data

**Supplementary Data 1**: Candidate genes based on genomic cross-referencing. List of candidate genes based on cross-referencing of PoolSeq, GWA, and Cline datasets.

**Supplementary Data 2:** Comprehensive annotation of the *A. distichus* neo-Y-linked genomic landscape. Comprehensive annotation and functional summary of all identified neo-Y-linked genes and open reading frames (ORFs) in *A. distichus*. This broad genomic summary includes strict gametologs, multi-chromosomal amplicons, neo-Y-specific duplications, and putative pseudogenes. Functional classifications, copy number status (including autosomal paralogs), and evolutionary classes are provided to characterize the complex genetic architecture, high repeat content, and degradation landscape of the *A. distichus* Y chromosome.

**Supplementary Data 3:** Strict X-Y gametologs identified on the *A. distichus* Y chromosome. Summary of strictly retained X-Y gametologs identified on the *A. distichus* Y chromosome. The table details the genomic coordinates of the neo-Y-linked genes and their corresponding X-linked counterparts (gametologs). Additional annotations include functional gene names, orthogroup classifications, and computational assessments of structural degradation and sequence completeness, providing insights into the evolutionary preservation of these ancestral sex-linked loci.

**Supplementary Data 4:** Candidate neo-Y-linked genes associated with sex determination, dimorphism, and color patterning. List of candidate neo-Y-linked genes associated with sex determination, sexual dimorphism, and color patterning.

